# The energy metabolism of *Cupriavidus necator* in different trophic conditions

**DOI:** 10.1101/2024.02.26.582058

**Authors:** Michael Jahn, Nick Crang, Arvid H. Gynnå, Deria Kabova, Stefan Frielingsdorf, Oliver Lenz, Emmanuelle Charpentier, Elton P. Hudson

## Abstract

The ‘knallgas’ bacterium *Cupriavidus necator* is attracting interest for biotechnological applications due to its extremely versatile metabolism. *C. necator* can use hydrogen or formic acid as an energy source, fixes CO_2_ *via* the Calvin-Benson-Bassham (CBB) cycle, and also grows well on various other organic acids and sugars. Its tri-partite genome is notable for its size and manifold duplications of key genes (CBB cycle, hydrogenases, nitrate reductases). Comparatively little is known about which of these (iso-) enzymes and their cofactors are actually utilized for growth on different energy sources.

Here, we investigated the energy metabolism of *C. necator* H16 by growing a barcoded transposon knockout library on various substrates including succinate, fructose, hydrogen (H_2_/CO_2_) and formic acid. The fitness contribution of each gene was determined from enrichment or depletion of the corresponding mutants. Fitness analysis revealed that 1) some, but not all, molybdenum cofactor biosynthesis genes were essential for growth on formate and nitrate respiration. 2) Soluble formate dehydrogenase (FDH) was the dominant enzyme for formate oxidation, not membrane-bound FDH. 3) For hydrogenases, both soluble and membrane-bound enzymes were beneficial for lithoautotrophic growth. 4) Of the six terminal respiratory complexes in *C. necator* H16, only some are utilized and utilization depends on the energy source. 5) Deletion of hydrogenase-related genes boosted heterotrophic growth, and we show that the relief from associated protein cost is responsible for this phenomenon.

This study evaluates the contribution of each of *C. necator*’s genes to fitness in biotechnologically relevant growth regimes. Our results illustrate the genomic redundancy of this generalist bacterium, and may inspire future strategies for strain engineering.

**Graphical Abstract:** 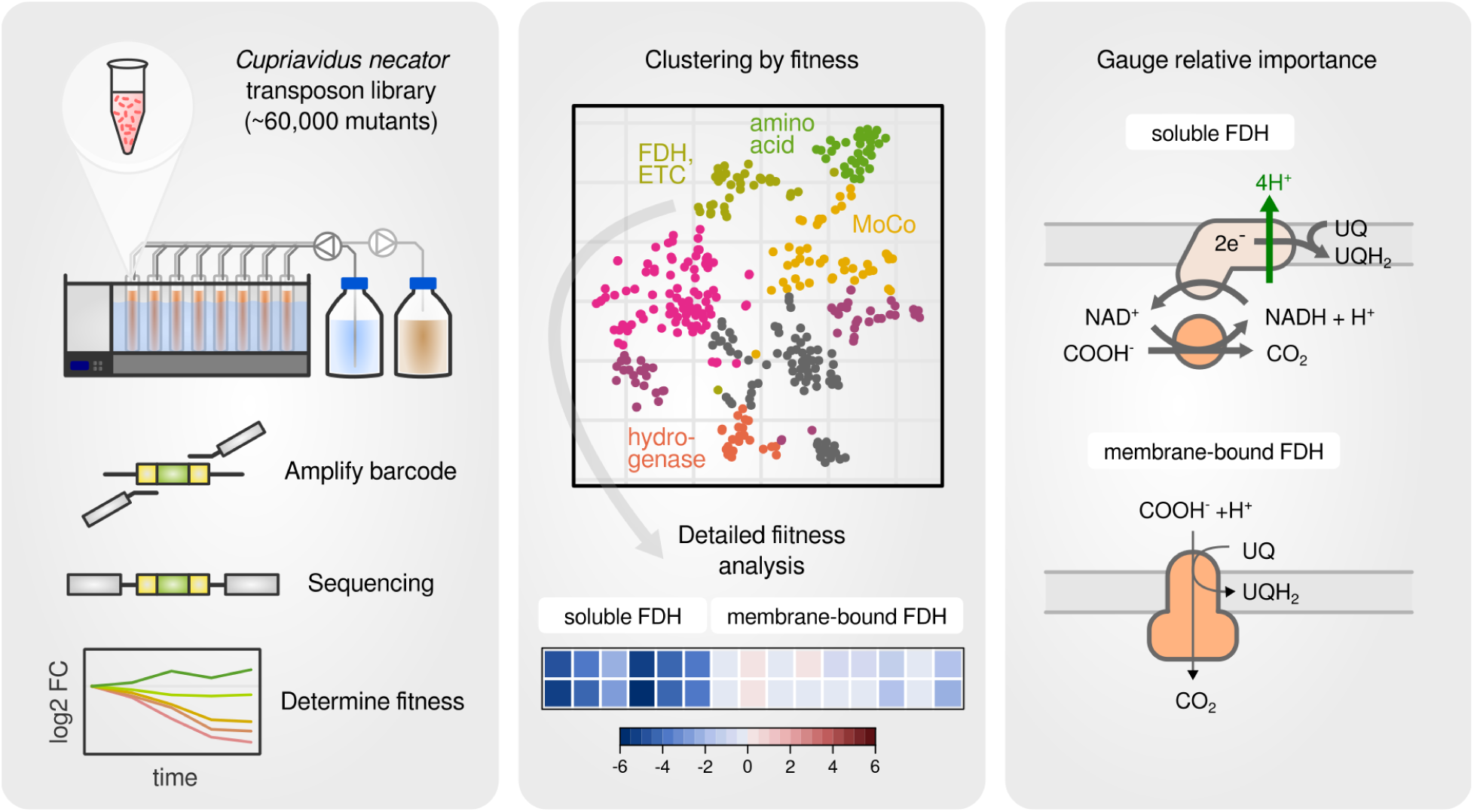

**Highlights:**

- a barcoded transposon library was used to assess gene fitness for hydrogenase, formate dehydrogenase and electron transport chain complexes
- the utilization of terminal respiratory complexes is substrate specific
- soluble formate dehydrogenase is more important than the membrane-bound enzyme
- soluble hydrogenase and membrane-bound hydrogenase are equally utilized
- inactivation of hydrogenase and its accessory genes leads to faster heterotrophic growth
- we demonstrate that faster growth is caused by the relief of protein synthesis burden
- fitness data is available in an interactive app at https://m-jahn.shinyapps.io/ShinyLib/

## Introduction

The soil bacterium *Cupriavidus necator* H16 (formerly *Ralstonia eutropha* H16) is a widely recognized model organism for chemolithoautotrophic growth (Bowien & Kusian, 2002, Pohlmann et al., 2006) and for the production of the bioplastic polymer polyhydroxybutyrate (PHB) (Reinecke & Steinbüchel, 2009, Panich et al., 2021). Its large inventory of enzymes enables it to grow on a wide range of heterotrophic substrates as well as on mixtures of molecular oxygen (O_2_), molecular hydrogen (H_2_), and carbon dioxide (CO_2_) (Cramm, 2009, Park et al., 2011, Pearcy et al., 2022). To fuel CO_2_ fixation by the Calvin Benson Bassham (CBB) cycle, *C. necator* generates ATP and reduction equivalents from the oxidation of molecular hydrogen (H_2_ → 2e^-^ + 2H^+^) or formic acid (HCOOH → CO_2_ + 2e^-^ + 2H^+^). Genome redundancy and phylogenetic comparison suggest that the ability for autotrophic growth was only recently acquired (Fricke et al., 2009). The large genome (∼6,600 genes) is divided into two chromosomes and one significantly smaller megaplasmid (pHG1) (Pohlmann et al., 2006). However, pHG1 encodes most of the proteins associated with lithoautotrophic growth and carries operons that extend metabolic versatility, such as nitrate respiration and aromatics degradation functions (Schwartz et al., 2003). Both pHG1 and chromosome 2 were likely acquired from other *Burkholderia* species by horizontal gene transfer, while chromosome 1 represents the evolutionarily most ancient DNA molecule (Fricke et al., 2009). In a recent study on the *C. necator* proteome we have shown that about 80% of its protein mass is encoded by chromosome 1 irrespective of the growth condition (Jahn et al., 2021). We have also shown that the expression of autotrophic pathways during heterotrophic growth leads to suboptimal utilization of resources. One example is the costly expression of CBB cycle genes during glycolysis, which leads to reassimilation of emitted CO_2_ (Shimizu et al., 2015, Subagyo et al., 2021), a process that is unlikely to have a biomass yield or growth rate benefit (Jahn et al., 2021). Another example is the redundancy of CBB cycle enzymes, which are encoded by two largely identical *cbb* operons on pHG1 and chromosome 2. Knocking out genes from either copy had no negative effect on autotrophic growth. The second *cbb* copy is therefore completely dispensable (Windhövel & Bowien, 1990, Jahn et al., 2021).

What remains less explored is *C. necator*’s energy metabolism reviewed by Cramm, 2009. It prefers organic acids over sugars (degraded *via* the Entner-Doudoroff pathway) to produce ATP during heterotrophic growth under respiratory conditions. The bacterium is also able to obtain energy from hydrogen oxidation *via* two remarkable O_2_-tolerant metalloenzymes, a soluble NAD^+^-reducing and a membrane-bound [NiFe]-hydrogenase (SH and MBH, respectively) (Lenz et al., 2018). The maturation of hydrogenases requires dedicated accessory proteins (encoded by the *hyp* genes) that synthesize and insert the Ni-Fe-(CN)_2_-CO cofactor into the enzymes (Böck et al., 2006, Bürstel et al., 2012, Schulz et al., 2019). The enzymes that accomplish formate oxidation, two soluble, NAD^+^-reducing (SFDH) and two membrane-bound formate dehydrogenases (MFDH), also require accessory proteins for maturation (Burgdorf et al., 2001, Cramm, 2008) and biosynthesis of the important molybdenum cofactor (MoCo) (Maia et al., 2015, Hille et al, 2020).

The soluble (de-)hydrogenases transfer electrons directly to NAD+ generating reducing power in the form of NADH. Membrane-bound (de-)hydrogenases couple the oxidation reaction to the reduction of a universal quinone e^-^ carrier which in turn drives the electron transport chain to build up a proton motive force. The electron transport chain of *C. necator* is highly complex (Cramm, 2008). Whereas *E. coli* has a total of three respiratory complexes, *C. necator* has no fewer than nine (three copies of *bo_3_*quinol oxidase, two copies of *bd* quinol oxidase, a *bc_1_*cytochrome reductase, and three different terminal cytochrome oxidases, *bb_3_*, *cbb_3_*, and *aa_3_*) (Cramm, 2008). These complexes are named after the different types of heme cofactors they contain and differ in their biochemical properties such as a) type of the e^-^ donor and acceptor, b) high or low affinity for the terminal e^-^ acceptor, and c) the number of protons pumped across the cytoplasmic membrane. Little is known about the utilization of these complexes in different trophic conditions. In addition to using oxygen as the terminal e^-^ acceptor, *C. necator* also has the ability to respire nitrate and other nitrogen oxides in anaerobic conditions. The genes encoding the four required enzyme complexes (nitrate, nitrite, nitric oxide, and nitrous oxide reductase) are located on pHG1 and expressed only when oxygen is limiting (Schwartz et al. 2003, Kohlmann et al., 2014).

In this study, we investigated which enzymes of energy metabolism contribute most to cell fitness in different growth regimes including heterotrophic, lithoautotrophic and formatotrophic growth. We focused on two areas of energy metabolism, first, the enzymes that regenerate reduced cofactors (NADH, quinol), and second, the electron transport chain that consumes the reduced cofactors to generate ATP. We wondered which of the different FDH and hydrogenase genes/operons are most utilized in *C. necator*. Which of the many accessory genes for FDH/hydrogenase maturation are truly required to produce functional enzymes? And what are the fitness costs associated with expression of these pathways? To answer these questions, we employed a barcoded transposon library (Wetmore et al., 2015, Price et al., 2018) that was recently used to study the versatile carbon metabolism of *C. necator* (Jahn et al., 2021). The mutant library was created by conjugating *C. necator* H16 with an *E. coli* donor strain, yielding around 60,000 individually barcoded transposon integration mutants (Figure 1 A, Wetmore et al., 2015). The barcodes were mapped to their respective genome integration site using the TnSeq workflow (Barquist et al., 2016) (Figure 1 A). The enrichment or depletion of individual mutants was then tracked by next generation sequencing (NGS) of the 20 nt barcode (Figure 1 B). We calculated a gene-wise fitness score from mutant abundances that represents the relative importance of a gene for a particular growth condition. The use of barcodes significantly increases the experimental throughput, which allowed us to probe a wide range of genes important for the various energy-generating pathways of *C. necator*.

**Figure 1:**
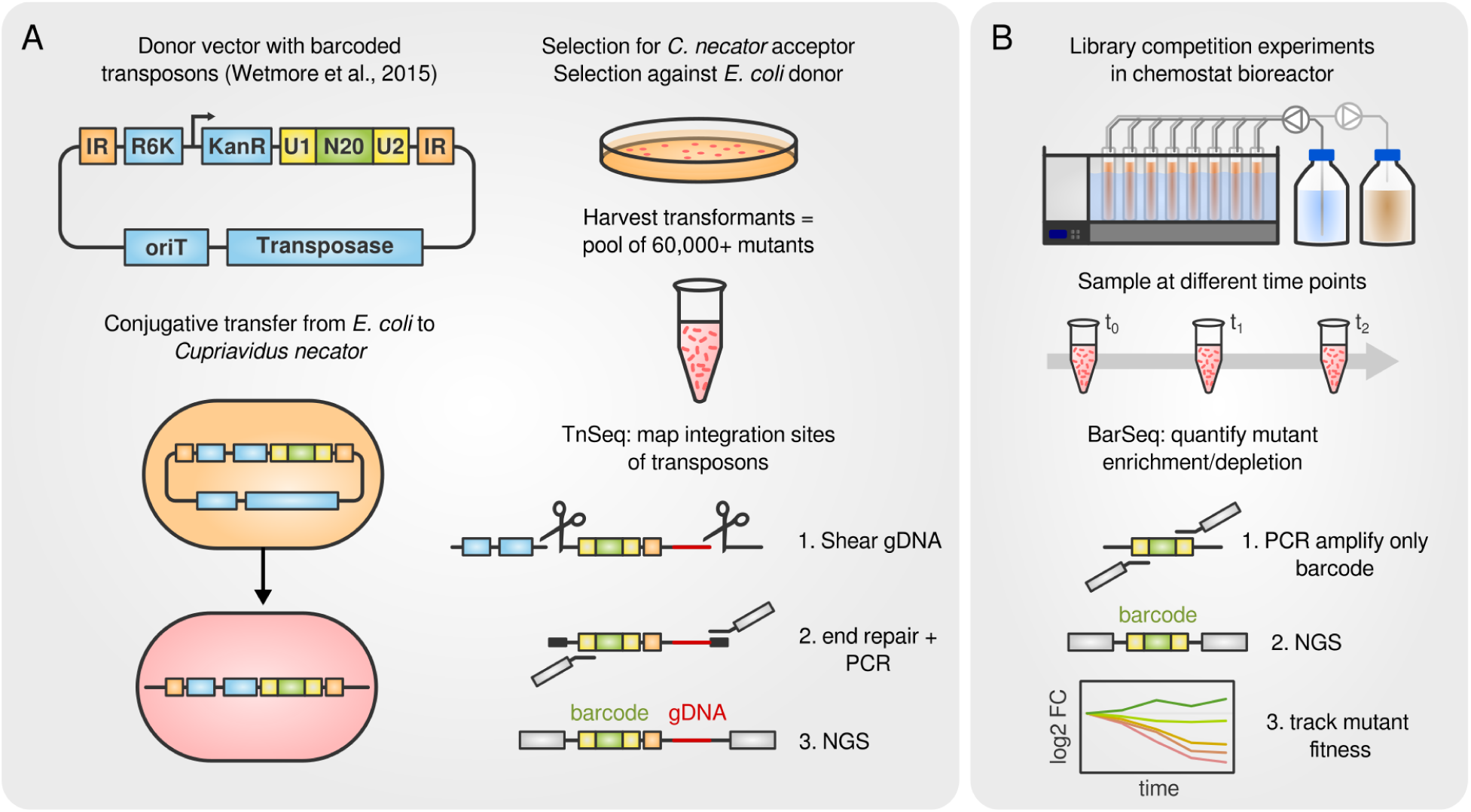
Creation of the *C. necator* transposon library and screening workflow. **A)** *C. necator* H16 was conjugated with an *E. coli* donor strain carrying a transposon integration construct (sketch of the vector was adopted from Wetmore et al., 2015). The construct contains two inverted repeat (IR) elements that serve as recognition sites for the transposase, an R6K plasmid replication origin, a kanamycin resistance cassette (kanR), two universal priming sites (U1, U2) and the 20 nt random barcode (N20). Transposon integration sites were mapped to barcodes using the TnSeq workflow. Genomic DNA was isolated, shear-fragmented and end-repaired. Transposon containing fragments were PCR amplified including the barcode and a portion of genomic DNA, and then subjected to NGS. **B)** Library competition experiments were mostly performed in chemostat bioreactors. Samples were taken after 0, 8, and 16 generations, genomic DNA was extracted and barcodes were sequenced using NGS. The changes in barcode abundance over time were used to calculate a fitness score for each gene.

## Results

### Genes involved in energy metabolism dominate fitness screening

We probed the contribution of genes to fitness by growing the transposon mutant library in five different trophic conditions: With fructose, succinate, or formate as carbon and energy source, and O_2_ as the terminal e^-^ acceptor, whereby formate is first oxidized to CO_2_ which is then fixed *via* the CBB cycle. In addition, we grew the mutant library in anoxic conditions with fructose as carbon and energy source and nitrate as the terminal electron acceptor (nitrate respiration), and finally, in the presence of a gas mixture of CO_2_, H_2_, and O_2_ (lithoautotrophy). Except for the latter, all cultivations were performed with two different feeding regimes, continuous feed (C) and pulsed feed (P), in order to select either for an advantage of growth rate or for biomass yield, respectively (Wides & Milo, 2018). Depletion or enrichment of mutants was tracked by barcode abundance and a fitness score calculated for each gene and growth condition (Figure 1B). Our analysis revealed 354 genes with a fitness score of *f* ≤ -2 or *f* ≥ 2 in any condition, which we clustered by similarity (Figure 2 A, Figure S1). The seven emerging clusters were each dominated by genes related to specific functional groups: Genes in cluster 1 were almost exclusively related to amino acid biosynthesis and depleted in all growth conditions (Figure 2 B,C). These genes became essential when the mutant library was transferred from the complete medium of pre-cultures to the minimal medium in bioreactor cultures. Genes in cluster 2 were depleted in formatotrophic growth and strongly enriched for formate dehydrogenases (assigned to 1-carbon metabolism pathways like ‘methane’, Figure 2 C) and electron transport complexes. Cluster 3 contained genes related to molybdenum cofactor biogenesis, and mutants were strongly depleted during nitrate respiration. The smallest cluster, number 4, contained almost exclusively genes related to H_2_ metabolism, and its mutants were strongly depleted during lithoautotrophic growth but, surprisingly, enriched in all other conditions. The remaining clusters 5 to 7 were weaker in terms of differential fitness and more heterogeneous in terms of assigned functions (Figure 2 C). We therefore focused our analysis on the three energy conversion-related groups formate dehydrogenases, hydrogenases, and the electron transport chain.

**Figure 2:**
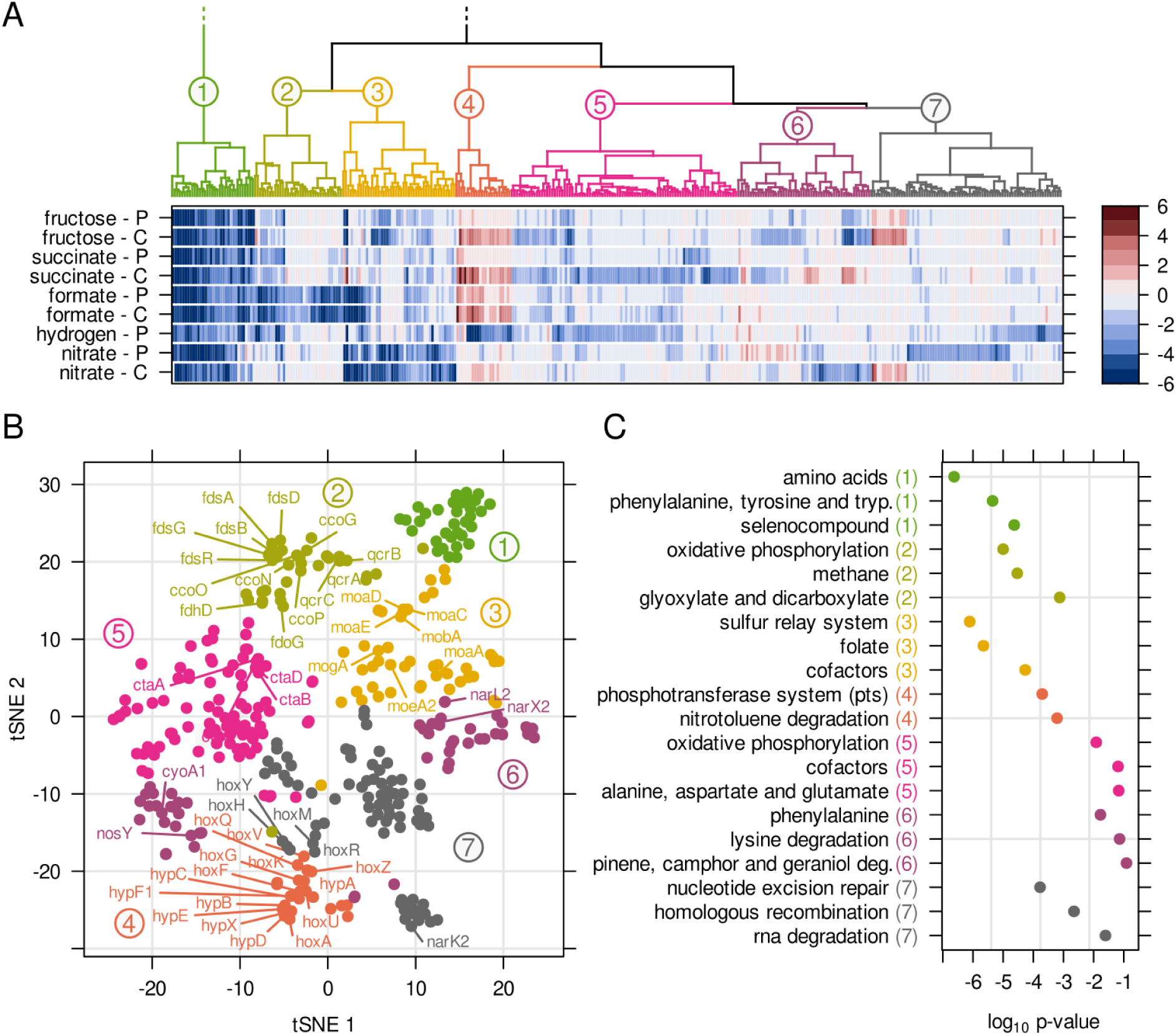
Clustering of genes with significantly changed fitness. **A)** Fitness scores for 354 genes exceeding a threshold *f* ≤ -2 or *f* ≥ 2. Genes were clustered based on similarity of fitness scores for the various growth conditions (dendrogram colored by cluster number). A cluster number of 7 was used after testing cluster separation by silhouette width, see Figure S1. P - pulsed feeding, C - continuous feeding. **B)** t-SNE dimensionality reduction of genes, color coded and labeled according to the clusters in A). **C)** Gene enrichment for KEGG pathways. The top three pathways with a p-value below 1 were selected. Colors and numbers in brackets correspond to the clusters in A).

### Fitness related to formate dehydrogenases and their cofactors

The formate dehydrogenases and nitrate reductases of *C. necator* are metalloenzymes that require a molybdenum cofactor (MoCo) which is synthesized by a dedicated pathway (Maia et al., 2015). The MoCo molecule is a polycyclic aromatic compound, molybdopterin, which binds a single molybdate anion (MoO_4_ ^2-^) *via* a dithiolene group (Iobbi-Nivol & Leimkühler, 2013). Four different steps are necessary to synthesize and insert MoCo into the apo-enzyme (Iobbi-Nivol & Leimkühler, 2013). First, the molybdopterin backbone is synthesized by cyclization of GTP to cGTP catalyzed by MoaA, conversion to pyranopterin by MoaC, and insertion of two sulfur residues by MoaD and MoaE (Figure 3 A). Only the genes *moaD*, moa*E* and the MoCo-unrelated fluoride transporter *moaF* are organized in an operon structure on chromosome 1 (Burgdorf et al., 2001). The *moaA* gene is located further downstream on chromosome 1 together with *mobA* and *moeA2*. We found that the *moaACDE* genes showed extremely low fitness scores and hence high importance for growth on formate and for nitrate respiration (Figure 3 A). Interestingly, *moaA* (H16_A2581) was only essential for nitrate respiration, but not for formatotrophic growth. A second non-quantified gene copy, *moaA2* (H16_B1466), probably compensates for the *moaA* knockout, suggesting that *moaA* is specific for GTP cyclization during nitrate respiration. The next step in MoCo biosynthesis is the import of the molybdate anion. In *C. necator,* a dedicated high affinity membrane transport system encoded by *modABC* exists, but its knock-out had no effect on fitness (Figure 3 A). An explanation for this might be the comparatively high concentration of MoO_4_ ^2-^ in the growth medium, permitting uptake by other, non-specific transporters (Xia et al., 2018). The molybdate anion is then inserted into molybdopterin by MogA and MoeA (Burgdorf et al., 2001), whereby MogA increases the affinity for molybdate at low concentrations without being strictly required (Iobbi-Nivol & Leimkühler, 2013). This was also reflected in the fitness scores: Both *mogA* as well as *moeA1* and *moeA2* knockouts showed slightly but not dramatically reduced fitness, confirming the non-essentiality of *mogA* and suggesting that *moeA2* can compensate for *moeA1* knock-out and *vice versa*. Knock-out of a third *moeA* copy, *moeA3*, showed no effect on fitness. Our results correspond to a previous report stating that *moeA1* becomes only essential when the other two *moeA* copies are inactivated (Burgdorf et al., 2001). Finally, the basic MoCo can be further modified by the MobA catalyzed attachment of GMP residues. The very low fitness score of *mobA* suggests that the GMP modification is strictly essential for FDH and nitrate reductase activity, a feature known from related *E. coli* enzymes of the DMSO reductase family to which *C. necator*’s FDHs belong (Iobbi-Nivol & Leimkühler, 2013). MobB, whose role in MoCo biosynthesis remains elusive, did not show any fitness effect for its three paralogs *mobB*, *mobB2* and *mobB3*.

**Figure 3:**
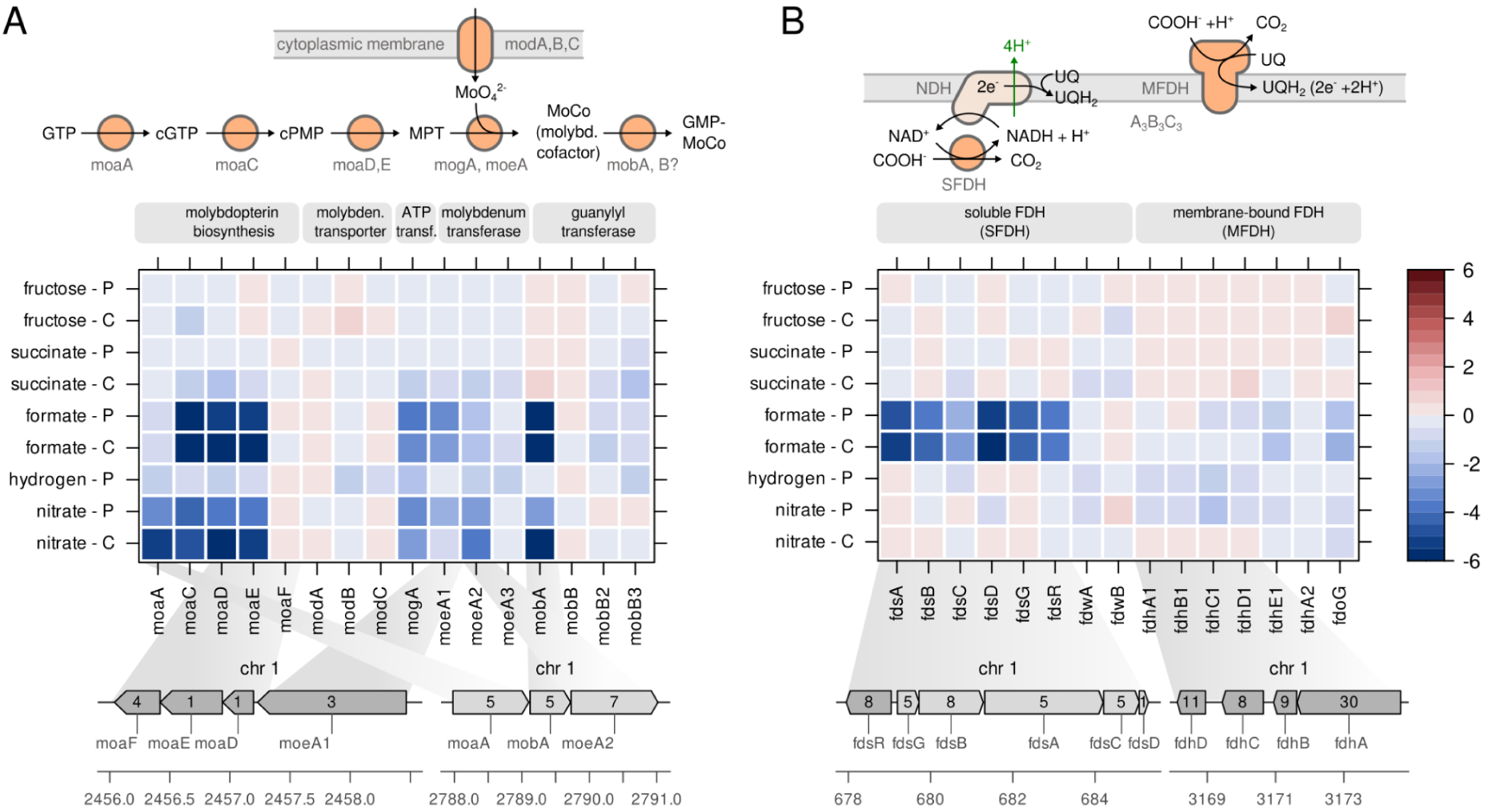
Fitness related to formate dehydrogenases and their cofactors. **A)** Fitness score for transposon insertion mutants of the molybdenum cofactor (MoCo) biosynthesis pathway. **B)** Fitness score for transposon insertion mutants of various formate dehydrogenase genes. A detailed description can be found in the Results section. The fitness of the *fdoHI* genes was not quantified. The scale bar indicates the gene fitness score after 8 generations of growth. Inset numbers in genome plots represent the number of transposon insertions per gene. cGTP - cyclic GTP, cPMP - cyclic pyranopterin monophosphate, MPT - molybdopterin, MoCo - molybdopterin cofactor, GMP-MoCo - GMP modified MoCo, SFDH - soluble formate dehydrogenase, MFDH - membrane-bound formate dehydrogenase.

The genome of *C. necator* possesses four known operons encoding FDH subunits. The *fdsGBACD* operon encodes the subunits of a soluble, heteromultimeric NAD^+^-reducing FDH (SFDH, Cramm 2008). The *fdsGBA* and *D* mutants showed greatly reduced fitness in formatotrophic growth (Figure 3 B). FdsA, FdsB and FdsG contain iron-sulfur clusters and FdsB also the molybdenum cofactor, which makes them likely participating in electron transfer. FdsD does not contain any such cofactor but may assist in maintaining the quaternary structure of the enzyme (Oh & Bowien, 1998, Radon et al., 2020). The role of FdsC remains unclear,but it was suggested that it transfers sulfur to MoCo or protects it from oxidation, prior to insertion into SFDH (Böhmer et al., 2014). The *fdsC* mutant showed moderately reduced fitness, suggesting that it is not strictly essential for growth on formate. However, since the essential *fdsD* gene is located downstream of *fdsC*, the reduced fitness for *fdsC* may also be caused by a negative effect on *fdsD* expression. Another soluble FDH (*fdwAB*) and two other membrane-bound formate dehydrogenases (MFDH) have been annotated but remain poorly functionally characterized (Burgdorf et al., 2001). The primary MFDH is encoded by the *fdhABCDE* operon; a secondary three-subunit enzyme is presumably encoded by the *fdoGHI* operon (Cramm 2008). While the *fsd* and *fdh* operons are located on chromosome 1, the *fdw* and *fdo* operons are located on chromosome 2, which is thought to have been acquired more recently during evolution (Fricke et al., 2009, Jahn et al., 2021). None of the *fdw*, *fdh*, and *fdo* knockouts showed a significant effect on fitness (Figure 3 B), suggesting that the SFDH encoded by the *fds* operon carries by far the main load of formate oxidation.

### Fitness related to hydrogenases and their cofactors

The [NiFe]-hydrogenases of *C. necator* are metallo-enzymes that require a nickel-iron-(CN)_2_-CO cofactor for the reversible oxidation of H_2_ into 2 e^-^ and 2 H^+^. All hydrogenase-related genes are located on the megaplasmid pHG1 (Schwartz et al., 2003). The metal cofactor is synthesized by a set of accessory Hyp proteins arranged in different operons on pHG1 (Figure 4 A, Fritsch et al., 2013). The largest operon, *hypA1B1F1C1D1E1X,* is located within an array of *hox* genes encoding the membrane-bound hydrogenase (MBH, *hoxKGZ*) and the regulatory hydrogenase (RH, *hoxABCJ*) (Schwartz et al., 2003). Two more partial copies of the *hyp* gene cluster can be found approximately 40 kb downstream as part of an operon including the soluble NAD^+^-reducing hydrogenase (SH, *hoxFUYHI*). The function of the *hyp* gene products is well-understood. HypX generates carbon monoxide (CO) from formyl-THF (Schulz et al., 2019), HypEF synthesizes the cyanide ligands from carbamoyl phosphate, HypCD serves as protein scaffold binding the central iron atom to which the CN and CO ligands are coordinated (Bürstel et al., 2012), and HypAB delivers the nickel ion for the active site. Fitness data for all *hyp* genes suggest that the primary, MBH-associated *hyp* operon is strictly essential for hydrogen assimilation (Figure 4 A), as knockout of these genes led to a dramatic decrease in lithoautotrophic growth. None of the alternative *hyp* genes were able to rescue a primary *hyp* gene knockout, nor did they have any effect on fitness. However, the dramatic decrease in fitness could also be explained by the disruption of *hoxA* transcription, which is located directly downstream of *hypX*. HoxA is the major transcriptional regulator of *hox*/*hyp* gene expression (see detailed discussion below). Moreover, we observed an unexpected increase in fitness for all primary *hyp* genes in non-lithoautotrophic conditions. The growth benefit was consistently greater with continuous cultivation (primarily growth rate selective), but still evident with pulsed feed cultivation (primarily yield selective).

**Figure 4:**
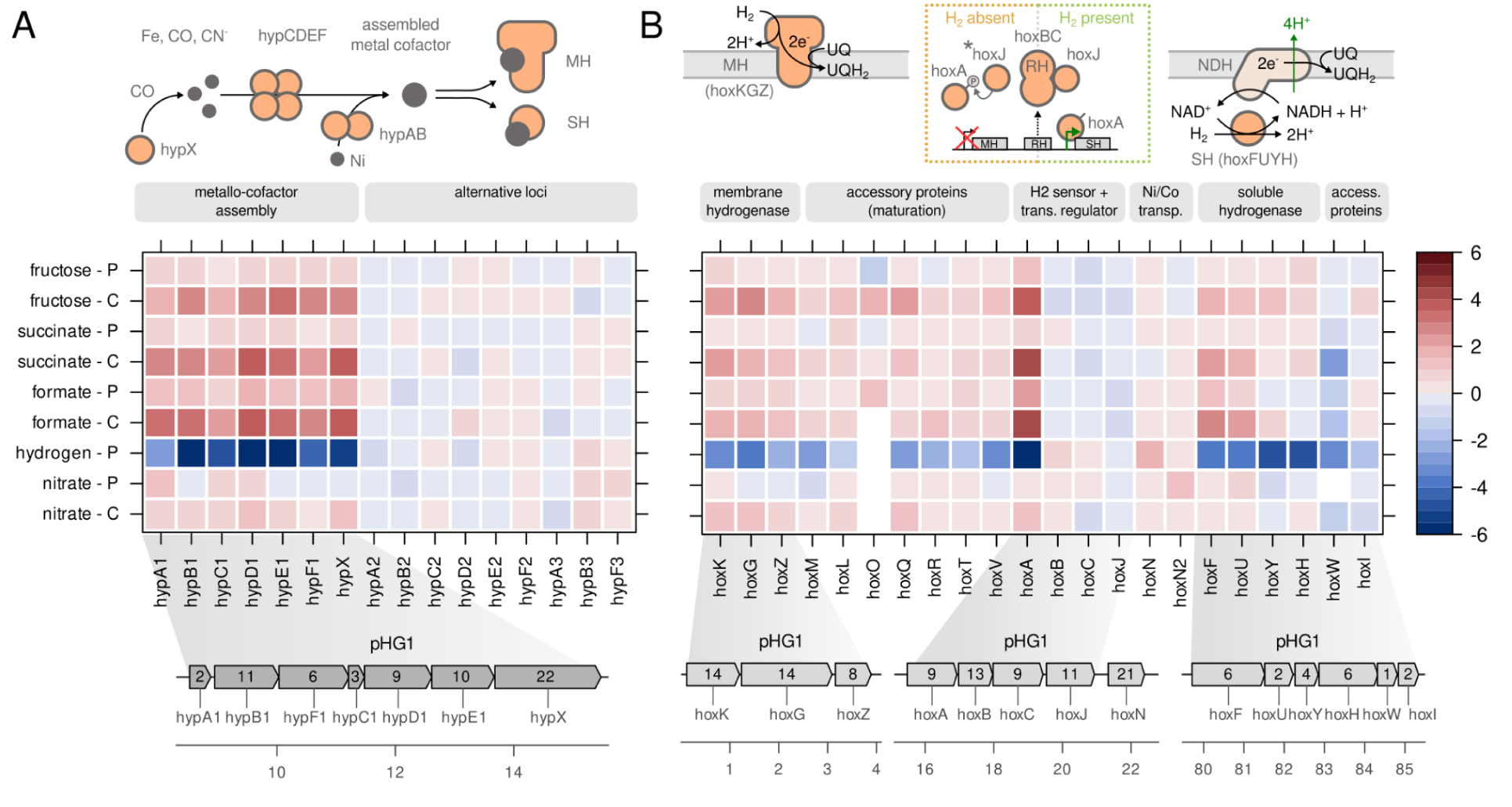
Fitness related to hydrogenases and their cofactors. **A)** Fitness score for the genes involved in biosynthesis of the Ni-Fe-(CN)_2_-CO cofactor (*hyp* genes). B) Fitness score for the genes forming membrane-bound hydrogenase (MBH), soluble hydrogenase (SH), regulatory hydrogenase (RH), and accessory genes for maturation of hydrogenases. For details, see the Results section. The scale bar indicates gene fitness scores after 8 generations of growth. Inset numbers in genome plots represent the number of transposon insertions per gene. Asterisk: Note that HoxJ is inactive in the H16 strain and can not regulate HoxA activity. CO - carbon monoxide, CN^-^ - cyanide, Ni - nickel, Fe - iron, UQ - ubiquinone, UQH_2_ - ubiquinol, NDH - NADH dehydrogenase (complex I).

The gene clusters encoding the two metabolic hydrogenases, MBH (*hoxKGZ*) and SH (*hoxFUYHI*), are separated by a 75 kb stretch containing accessory *hox* and *hyp* genes (Figure 4 B). Inactivation of *hoxKGZ* and *hoxFUYHI* genes led to decreased fitness in lithoautotrophic conditions, however, fitness was not as strongly decreased as for the *hyp* operon. This indicates that the inactivation of either MBH or SH can be partially compensated for by the activity of the respective other hydrogenase. Mutation of the accessory genes *hoxMLOQRTVW* responsible for hydrogenase maturation also showed reduced fitness during lithoautotrophic growth on a similar scale as that of the structural hydrogenase subunits. The genes for a third, non-enzymatic hydrogenase (*hoxBC*) termed the regulatory hydrogenase RH are located between the MBH and SH operons (Figure 4 B, Lenz et al., 2002). The RH is the hydrogen sensor of a two-component transcriptional regulation system. In the absence of H_2_, the histidine kinase HoxJ is phosphorylated and transfers its phosphate residue to HoxA, the master transcriptional regulator, turning it off. In the presence of H_2_, RH binds HoxJ and leaves the non-phosphorylated HoxA in its active state. The *C. necator* H16 strain used in this study lost the ability to respond to H_2_ due to a mutation in *hoxJ*, turning *hyp*/*hox* gene expression constantly on in energy-limited conditions (Lenz et al., 2002, Lenz et al., 2018). Accordingly, *hoxB*, *hoxC* and *hoxJ* genes were dispensable while the knockout of *hoxA* led to a dramatic loss of fitness in lithoautotrophic conditions. On the other hand, fitness of the *hoxA* mutant increased for all other growth conditions similar to *hyp* gene knockout, suggesting that unnecessary hydrogenase expression imposes high metabolic cost.

### Fitness related to electron transport chain complexes

1. *C. necator* H16 is strictly dependent on respiration in order to generate sufficient ATP to support growth. Electrons stripped from a wealth of substrates are either transferred directly (e.g. from MBH, MFDH) to universal electron carriers such as ubiquinone (UQ), or indirectly *via* NADH (e.g. from SH, SFDH) and the activity of NADH dehydrogenase (respiratory complex I) to UQ (Cramm, 2008). Electrons from UQ can then take different paths through the electron transport chain (ETC), generating a proton-motive force across the cytoplasmic membrane that is used to generate ATP through the rotational movement of ATP synthase. Judging by the number of different complexes potentially involved in electron transport, the ETC of *C. necator* is both highly flexible and highly complex (Figure 5). The structural and functional differences of ETC complexes are described in more detail elsewhere (Cramm, 2008, Morris & Schmidt, 2013).

Here we focus on basic functional properties such as electron acceptor affinity and contribution to the proton gradient. The direct route that electrons can take from the e^-^ donor ubiquinol (UQH_2_) to the terminal e^-^ acceptor oxygen is through the quinol oxidase complexes *bo_3_* and *bd*. In *C. necator*, the *bo_3_*complex is encoded by three different *cyoABC* operons and the *bd* complex by two *cydAB* operons (Figure 5). The *bd* family complexes do not pump protons, but nevertheless contribute to the membrane potential by consuming two protons through the reduction of O_2_ to H_2_O at the cytoplasmic side of the membrane (Morris & Schmidt, 2013). Of the *bo_3_* and *bd* related genes, none showed any significant effect on growth after inactivation. Although it is possible that duplicate genes compensate for the inactivation of the respective complexes, we assume that an overwhelming majority of ETC flux takes the alternative route *via* cytochrome reductase (*bc_1_* complex). *C. necator* has one copy of the *bc_1_* complex encoded by the *qcrABC* operon and inactivation of any of its genes led to significantly reduced fitness in all conditions except with nitrate as the e^-^ acceptor (Figure 5). The fitness penalty was highest for formatotrophy and lowest for growth on fructose. The *bc_1_*complex, which is the most abundant ETC complex in *C. necator* (Figure S2), transfers two e^-^ from UQH_2_ to a cytochrome *c* carrier protein and pumps a total of four protons to the periplasmic side.

**Figure 5:**
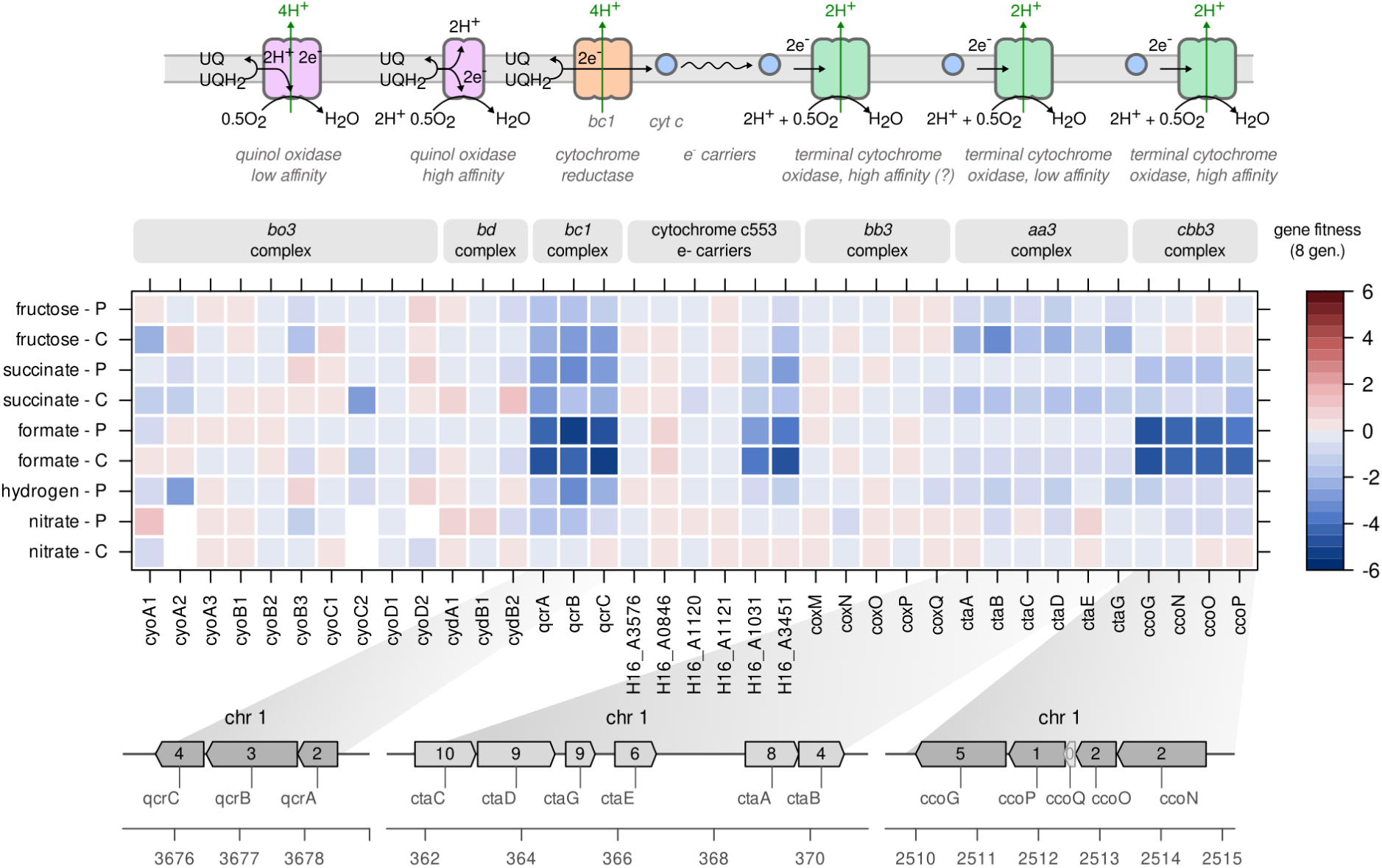
Fitness related to electron transport chain complexes. Fitness score for the genes encoding subunits of different respiratory complexes in the electron transport chain. For details, see the Results section. The scale bar indicates gene fitness scores after 8 generations of growth. Inset numbers in genome plots represent the number of transposon insertions per gene. UQ - ubiquinone, UQH_2_ - ubiquinol, e^-^ - electrons.

Cytochrome *c* is then oxidized by one of three complexes, *bb_3_*, *aa_3_* or *cbb_3_*, all of which transfer the e^-^ to O_2_ to reduce it to water. Inactivation of the genes encoding the *bb_3_* complex (*coxMNOPQ*) did not show any effect on fitness. Inactivation of the genes encoding the *aa_3_*complex (*ctaABCDEG*) on the other hand led to moderately reduced fitness in fructose and succinate driven growth, predominantly with pulsed feeding. The strongest effect on fitness was observed for the genes of the *cbb_3_*complex (*ccoGNOP*) during growth on formate, to a lesser extent for succinate, and not at all for fructose. This suggests that *C. necator* prefers the longer but more energy-efficient route *via* cytochrome reductase and oxidase (4+2 protons pumped per UQH_2_), instead of the direct reduction of O_2_ through quinol oxidases (2/4 protons pumped per UQH_2_). The low O_2_ affinity complex *aa_3_* was preferred in heterotrophic growth and the high O_2_ affinity complex *cbb_3_* in formatotrophic growth. This result is supported by mass spectrometry data, where the most abundant *aa_3_* subunit (CtaC) was more abundant in fructose, and the most abundant *cbb_3_* subunit (CcoO) more abundant in the formate condition (Figure S2).

1. *C. necator* also has the ability to respire anaerobically using nitrate as the terminal electron acceptor. Cells were cultivated under anaerobic conditions (Figure S3) with fructose or formate as carbon and energy sources (Figure S4). We found that *C. necator* was able to grow on fructose with nitrate supplementation, but remarkably, growth on formate and nitrate was impossible (Figure S4B). Next, we analyzed the effect on fitness from knock-out of the four denitrification pathways using our transposon mutant library. The *C. necator* genome contains operons for enzymes catalyzing all four steps of denitrification from nitrate (NO_3_^-^) *via* nitrite (NO_2_^-^), nitric oxide (NO) and nitrous oxide (N_2_O) to dinitrogen (N_2_), all located on chromosome 2 or pHG1 (Cramm, 2008). Surprisingly, we found no change in fitness for denitrification related genes (Figure S5), regardless whether nitrate was respired or not. The only two exceptions were the regulatory genes *narX2* and *narL2*, whose mutants had slightly reduced fitness in one nitrate respiration condition. This result is puzzling as it is known that the *nar*, *nir*, *nor* and *nos* operons are strongly upregulated during nitrate respiration, and that the intermediary products of each denitrification step can be detected in *C. necator* (Kohlmann et al., 2014). We assume that gene knockouts were compensated for by iso-enzymes of the denitrification pathway, thus rescuing the mutants.

### Protein cost explains growth advantage of hydrogenase mutants

A surprising result was the positive fitness score of a range of hydrogenase mutants (Figure 4). We estimated the growth advantage of *hox* and *hyp* mutants based on the change of NGS read counts over time (Figure 6 A, Methods). We found that inactivation of the accessory *hypABCDE* and *hypX* genes as well as inactivation of the master regulator *hoxA* led to an estimated increase in growth rate from 0.1 h^-1^ for the average population to 0.12-0.14 h^-1^ for the mutants. Such a pronounced increase in growth rate implies the removal of a considerable burden associated with hydrogenase-related genes. We therefore investigated two scenarios, the protein cost hypothesis and the metabolic cost hypothesis, which could explain this observation.

**Figure 6:**
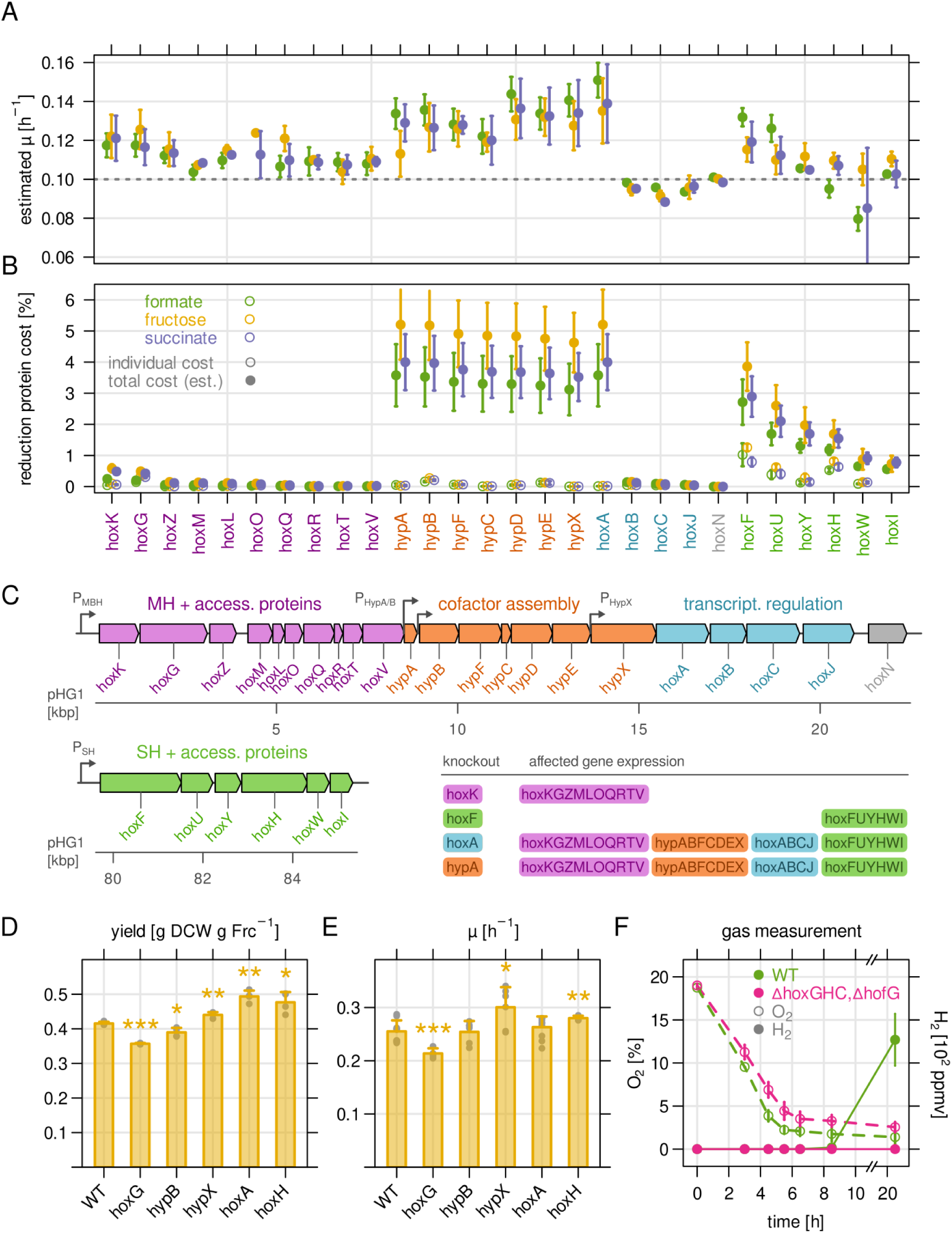
Protein cost explains growth advantage of hydrogenase mutants. **A)** Estimated growth rate µ of hydrogenase-related transposon mutants. The growth rate was calculated from the log_2_ fold change in mutant abundance over time compared to the population average. The dashed line marks the nominal growth rate for bioreactor experiments. **B)** Corresponding protein cost for the knockout mutants in A) in percent of the total protein mass. Open symbols, individual cost. Closed symbols, protein cost when disruption of downstream gene expression is taken into account. **C)** Organization of the various hydrogenase-related genes. Membrane-bound hydrogenase MBH (purple), accessory *hyp* genes (orange), and regulatory *hox* genes (blue) are located sequentially on megaplasmid pHG1. Genes for soluble hydrogenase (SH) are located around 60 kb downstream of the MBH operon. Expression of the promoters P_MBH_ and P_SH_ is controlled by the HoxA master regulator. Inset table: Effect on transcription of the selected gene knockouts. It is assumed that knockout of any of the primary *hyp* genes (orange) also disrupts *hoxA* expression. **D)** Biomass yield of selected, in-frame knockout mutants for batch cultures in fructose-supplemented minimal medium. Stars indicate the significance level of a two-sided t-test of each mutant against wild type. *, p ≤ 0.05. **, p ≤ 0.01. ***, p ≤ 0.001. **E)** Growth rate determined from optical density measurement for the same cultures as in D). **F)** Gas chromatography measurement in the headspace of batch cultures of wild-type *C. necator* H16 (green), and a mutant lacking all known hydrogenases (Δ*hoxG*, Δ*hoxH*, Δ*hoxC* and Δ*hofG*, in red). Open symbols, O_2_ measurement. Closed symbols, H_2_ measurement. All subfigures: Points and bars represent the mean of at least four biological replicates except F) where three replicates were used. Error bars represent standard deviation.

The protein cost hypothesis states that a bacterium is fundamentally limited in its capacity to synthesize proteins (Molenaar 2009, Jahn et al., 2018, Zavrel et al., 2019, Jahn et al., 2021). As a consequence, the utilization of proteins becomes important to optimize growth rate, and every idle enzyme constitutes an additional cost. The knockout of an unutilized enzyme leads to a reduction of the protein cost. In order to estimate the *individual* protein cost of hydrogenase related genes, we determined the protein mass fraction for each Hox/Hyp protein from mass spectrometry data acquired in a previous study (Figure 6 B, Jahn et al., 2021). We then summed up the protein cost for the inactivated gene and all downstream affected genes according to their genomic arrangement and known transcripts, yielding the *total* cost (Figure 6 B, C). For example, inactivation of *hoxK* leads to a low reduction of protein cost because it constitutes only the sum of the protein mass for the lowly expressed *hoxK* to *hoxV* genes. The inactivation of *hoxF* disrupts expression of all SH genes, yielding intermediate reduction of protein cost. Inactivation of *hoxA* on the other hand disrupts expression of all *hox* and *hyp* operons, resulting in a strong reduction of protein cost. Knockout of the primary *hyp* genes had the same effect as a *hoxA* knockout, from which we conclude that *hoxA* expression is dependent on the secondary *P_HypA_* or *P_HypB_* promoters (Lenz et al., 2002). The pattern of total protein cost reduction correlated well with the estimated increase in growth rate (Figure 6 A/B, Figure S6).

To validate the observed fitness advantage, we determined the biomass yield and growth rate of selected in-frame knockout mutants representative for MBH (*hoxG*), SH (*hoxH*), accessory *hyp* genes (*hypB*, *hypX*), and the master transcriptional regulator (*hoxA*). The mutants were grown in separate bioreactor batch cultures with fructose as the carbon source, such that hydrogenase expression constitutes a growth burden. Unlike transposon insertions, these knockouts are not expected to affect the expression of downstream genes. We therefore expected the highest increase in biomass yield or growth rate for the *hoxA* mutant, a moderate increase for *hoxH*, and little to no increase for the lowly expressed *hoxG*, *hypB* and *hypX* mutants. Indeed, the biomass yield was highest for the *hoxA* deletion, followed by *hoxH* and hypX, and lower than or equal to wild type for *hoxG* and *hypB* (Figure 6 D). The maximum growth rate determined from OD_720_ readings showed an overall similar result (Figure 6 E), with the exception of *hoxA*, which showed a lower growth rate than expected. However, we observed a shorter lag phase for *hoxA*, *hypB* and *hypX* mutants, which could also represent a growth advantage (Figure S7).

Finally, we tested the metabolic cost hypothesis. In heterotrophic conditions, hydrogenases could catalyze the reverse reaction of H_2_ oxidation – the formation of H_2_ from protons and an electron donor such as NADH (2H^+^ + 2e^-^ → H_2_), which could drain the NAD(P)H pool. The dissipation of excess reducing power in the form of NADH is for example known from other microbial phyla such as cyanobacteria (Tamagnini et al. 2002, Krishnan et al., 2018). It has been reported that *C. necator* produces H_2_ when the terminal electron acceptor, oxygen or one of the nitrogen oxides, is limiting and NADH accumulates (Kuhn et al., 1984). In order to determine if *C. necator* wastes reducing power through the action of hydrogenases, the wild-type H16 strain and a deletion mutant defective in all hydrogenase activity (Δ*hoxG*, Δ*hoxH*, Δ*hoxC* and Δ*hofG*) were cultured in batch with fructose and glycerol to maximize hydrogenase expression (Figure 6 F). No hydrogen evolution was detectable unless oxygen was severely limited. Hydrogen was only detected for the wild type strain at the last probed time point (20.5 h) at a concentration of around 1300 ppm (0.13% v/v). Such oxygen-depleted conditions were not present in our bioreactor experiments, ruling out the metabolic cost hypothesis and strengthening the protein cost hypothesis.

## Discussion

*C. necator* has attracted interest not only for its ability to produce the storage polymer PHB in high concentrations, but also for its extremely versatile carbon and energy metabolism. Many of the initial groundbreaking studies identifying the key enzymes for its remarkable metabolic capabilities date back to the 1980s and 1990s (Schneider & Schlegel, 1976, Friedrich et al., 1981, Friedebold & Bowien, 1993, Oh & Bowien 1998). Another important milestone was the publication of the full genome sequence (Pohlmann et al., 2006), which made it possible to compile a detailed list of all theoretically present energy acquisition pathways (Cramm, 2008). However, since then most studies on *C. necator* have focused on enhancing PHB production or engineering pathways for biosynthesis of other compounds replacing PHB (Kaddor et al., 2011, Tang et al., 2020, Bommareddy et al., 2020, Janasch et al., 2022). Our study, on the other hand, attempts to gain a holistic view of *C. necator*’s energy metabolism.

We have recently studied the carbon metabolism of *C. necator* from the perspective of resource allocation in different trophic conditions (Jahn et al., 2021). To this end, a barcoded transposon library was created to quantify gene essentiality in the central carbon metabolism, particularly the Calvin cycle which is used for CO_2_ fixation. Here, we exploited our transposon library to probe the energy metabolism of *C. necator*. Depletion or enrichment of individual strains thereby revealed the effect of inactivation of a gene on growth rate and biomass yield, which we summarize with the term ‘fitness contribution’. Since our data was particularly enriched for specific clusters of genes that affect fitness for formate and hydrogen assimilation as well as the molybdenum cofactor assembly (Figure 2 B), our study focused on these pathways.

The molybdenum cofactor (MoCo) is essential for the function of the two structurally and functionally related enzymes formate dehydrogenase and nitrate reductase (Cramm, 2008). Our results confirmed the essentiality of various enzymes responsible for MoCo biosynthesis. While most enzymes were essential for both formate assimilation and nitrate respiration, *moaA* was only essential for the latter. A second copy, *moaA2*, can probably compensate for this knockout specifically during formatotrophic growth. A set of genes responsible for molybdate insertion into molybdopterin (*mogA*, *moeA1* and *moeA2*) led to slightly reduced but not completely abolished growth. This confirms that *mogA* is beneficial but not essential (Iobbi-Nivol & Leimkühler, 2013), and that both *moeA* copies are functional and can partially compensate for the knockout of the respective other copy. A third copy, *moeA3*, was hypothesized to play the same role (Cramm, 2008), but our fitness data indicate that this copy is fully dispensable in the conditions tested.

The molybdenum cofactor is an essential component of *C. necator*’s two types of formate dehydrogenases, SFDH and MFDH. The molecular structure and enzymatic mechanism of the main SFDH (*fds* operon) have been studied in great detail (Friedebold et al., 1993, Niks et al., 2016, Hille et al., 2020). Much less is known about the two MFDH enzymes and the second copy of the SFDH, which share only limited similarity on gene level. It is unknown to which extent these formate dehydrogenases are active during formatotrophic growth and whether all or some of them carry metabolic flux. Fitness data from our library experiments clearly suggests that the SFDH encoded by the *fds* operon is the enzyme carrying the main metabolic flux of formate oxidation. All *fds* mutants had strongly reduced fitness while all other FDH mutants did not. In library screenings, we often see a partial decrease in fitness when multiple iso-enzymes with similar activity share the metabolic load (Miao et al., 2023), but the FDH genes did not show such a ‘division of labor’. This also means that the overwhelming majority of electron flux from formate oxidation is channeled towards NADH, and only indirectly towards ubiquinone/ubiquinol in the electron transport chain.

The fitness screening for hydrogenase genes –unlike formate dehydrogenase– suggests that both soluble and membrane-bound hydrogenase are equally utilized for hydrogen oxidation. Mutants for the SH genes *hoxFUYHI* showed moderately reduced fitness scores, similar to the fitness from *hoxKGZ*, while fitness scores for the *hoxA* knockout, which disrupts all *hox* gene expression, were around twice as low. The *C. necator* hydrogenases have been of interest for their oxygen tolerance, and have been structurally and functionally well characterized (Saggu et al., 2009, Fritsch et al., 2013). But less is known about their physiological relevance in different growth conditions. It has been shown that hydrogenase expression is upregulated when the preferred carbon source (e.g. fructose, succinate) is depleted in the culture medium and that expression levels of both hydrogenases are comparable (MBH around 60% of SH, Schwartz et al., 1998, Lenz et al., 2002, Jugder et al., 2015). But the physiological importance in terms of metabolic flux has not yet been quantified. Our results indicate that SH and MBH most likely have a similar contribution in carrying the metabolic flux from H_2_ to reduction equivalents. The SH and MBH enzymes require accessory proteins for biosynthesis of cofactors (*hyp* genes) and maturation (*hoxMLOQRTVW*). Of the three *hyp* operons on pHG1, only mutants of the primary *hyp1* operon showed a reduction in fitness. The strong phenotype was remarkably similar to that of the *hoxA* mutant, suggesting that the KO of these genes disrupts the downstream *hoxA* expression, and leads to complete loss of H_2_ oxidation capacity. The *hydrogenase* maturation genes were all essential for lithoautotrophic growth with the exception of *hoxN/N2*, which encode nickel transporters (Eitinger & Friedrich, 1991) that are most likely not important when metal concentration in the environment is sufficient.

In formatotrophic or lithoautotrophic lifestyle, redox power generated from formate or H_2_ oxidation can take two different routes to fuel the energy requirements of *C. necator*. The NADH from soluble FDH/SH can reduce ubiquinone (UQ) *via* the NADH dehydrogenase (NDH, also known as respiratory complex I). All subunits of NDH are fully essential, as the library contained no transposon mutants of the *nuo* operon encoding them. The second route is the direct e^-^ transfer to UQ. The reduced UQH_2_ can then be used as a substrate to pump protons by at least three ETC complexes. We found that *C. necator* prefers the route via cytochrome *c* reductase (*qcrABC*), which is 50% more energy efficient than quinol oxidases in terms of protons pumped (6 vs 2/4), but more complex and therefore likely associated with a higher protein cost. We also found that, of the three annotated terminal cytochrome *c* oxidases, the *cbb_3_*complex is preferred during formatotrophic growth and the *aa_3_* complex for growth on fructose. We note that ETC complexes have different affinity for oxygen, but we only cultivated the library with unlimited O_2_ supply (except for nitrate respiration). None of the complexes was clearly essential for lithoautotrophic growth.

When oxygen is replaced by nitrogen oxides, a completely different set of terminal oxidases is used for anaerobic respiration – the dedicated nitrate, nitrite, nitric and nitrous oxide reductase complexes (Cramm, 2008, Kohlmann et al., 2014). We grew the transposon library in denitrifying conditions and verified that nitrate supplementation was strictly required for growth. Yet, we found none of the annotated terminal reductases to be essential. A combination of the following two hypotheses could explain this surprising result: 1) Inactivation of individual denitrification genes can be compensated by iso-enzymes. For example, a knockout of the primary nitrate reductase (NAR) could be compensated for by the function of either of the two secondary reductases NAP and NAS, although the latter is most likely not expressed in presence of the nitrogen source we used, ammonium chloride (Warnecke-Eberz & Friedrich, 1993). 2) Nitrate was present at non-limiting concentration, such that *C. necator* was not dependent on the entire denitrification pathway from nitrate to N_2_, making downstream denitrification genes non-essential. However, the genes for biosynthesis of the molybdenum cofactor were strictly essential for denitrification (Figure 2 A). This cofactor is an essential part not only of FDH, but also of all three nitrate reductases (NAR, NAP, NAS), implying that one or more of these enzymes were actively used. We noted that nitrate respiration was impaired when cells were grown on formate instead of fructose (Figure S4). It has previously been reported that *C. necator* cannot respire nitrate when grown lithoautotrophically (Tiemeyer et al., 2007), and this seems to apply to formate as well. Our results did not provide any hints as to why litho- and formato-autotrophic growth is incompatible with nitrate respiration.

Fitness data for lithoautotrophic growth surprisingly revealed that inactivation of several hydrogenase-related genes boosted growth in heterotrophic conditions. We tested two hypotheses, hydrogenases causing metabolic cost by wasteful conversion of NADH to H_2_, or protein cost by production of enzymes (and cofactors). We found that the increase in growth rate correlated well with the estimated protein mass of hydrogenases and accessory proteins. Moreover, no emission of molecular hydrogen was detected in heterotrophic growth that would support the metabolic cost hypothesis. The expression of unutilized hydrogenases therefore constitutes a severe growth burden for *C. necator*, at least for strain H16, which was reported to be defective in H_2_-dependent regulation of *hox/hyp* gene expression (Lenz & Friedrich, 1998). A recent study identified targets for rational engineering of *C. necator* towards faster growth on formate, and reached similar conclusions (Calvey et al., 2023): Almost all of the spontaneous mutations inactivated either SH, MBH, or the CBB cycle enzymes, whose under-utilization we have described previously (Jahn et al., 2021). The complete removal of the pHG1 megaplasmid encoding all hydrogenases led to one of the biggest improvements in growth rate (Calvey et al., 2023). The increase in maximum growth rate of around 20% for ΔpHG1 corresponds well to the increase of yield or growth rate we observed for selected *hox*/*hyp* mutants, but even more so for the growth rate estimated from library experiments (Figure 6). Altogether, our results are in line with the findings from Calvey et al. and mount more evidence for the importance of protein cost in *C. necator*. The tight regulation of hydrogenase expression in strains with functional *hoxJ* signifies the high protein cost associated with litho-autotrophic lifestyle. The detrimental effect of a protein burden on bacterial growth is well-studied in model bacteria (Hui et al., 2015, O’Brien et al., 2016, Mori et al., 2017, Jahn et al., 2018). Microbes nevertheless employ the strategy of keeping enzyme reserves in order to cope with fluctuating environments, changing carbon/energy supply, and stress (Basan et al., 2016, Mori et al., 2017, Jahn et al., 2021). The genome of *C. necator* appears well adapted to such a lifestyle, but less suitable for biotechnological application. Our results open up further avenues for the rational engineering of *C. necator* for improved growth in biotechnologically relevant settings.

## Methods

### Strains

*Cupriavidus necator* H16 was obtained from the German Collection of Microorganisms and Cell Cultures, strain number DSM-428. Strains carrying individual in-frame deletions for *hoxG* (Bernhard et al., 1996), *hypB* (Dernedde et al., 1996), *hypX* (Buhrke and Friedrich, 1998), *hoxA* and *hoxH* (Bernhard et al., 1996) have been described previously. The negative control strain lacking all four hydrogenases (AH, MBH, RH, SH) was created by serial in-frame deletions of the respective hydrogenase subunit genes. *C. necator* HF364, which already carried deletions in the large subunit genes of the MBH, RH, and SH, served as starting point (Schäfer et al., 2013). In this strain, the native AH promoter was already replaced by the MBH promoter. A PCR fragment was amplified with primers 1 and 2 (Table S2), using the megaplasmid pHG1 as a template. The fragment contained the AH large subunit gene *hofG* as well as flanking regions. The fragment was digested with XbaI and subsequently ligated into XbaI-cut pLO1 (Lenz et al., 1994), resulting in the plasmid pGE815. Next, a 1131 bp fragment was removed from the *hofG* gene by restriction using SexAI and subsequent religation, resulting in plasmid pGE816. Double homologous recombination using the suicide vector pGE816 facilitated the incorporation of the *hofG* deletion into *C. necator* HF364, yielding the final Δ*hoxG*, Δ*hoxH*, Δ*hoxC*, Δ*hofG* deletion.

### Creation of barcoded *C. necator* transposon library

The creation of the barcoded transposon knockout library has been described in detail in Jahn et al., 2021 and Wetmore et al., 2015. Briefly, *C. necator* H16 wild-type was conjugated with an *E. coli* APA766 donor strain containing a barcoded transposon library. The strain is auxotrophic for DAP, the L-Lysin precursor 2,6-diamino-pimelate, to allow for counter selection. Overnight cultures of *E. coli* APA766 and *C. necator* H16 were prepared. The APA766 culture was supplemented with 0.4 mM DAP and 50 µg/mL kanamycin. Fresh cultures were prepared from precultures and then harvested during exponential growth phase by centrifugation for 10 min, 5,000 xg, RT. Supernatant was discarded, cell pellets were resuspended in residual liquid, transferred to 2 mL tubes, washed twice with 2 mL PBS, and finally resuspended in a total amount of 500 µL PBS. Cell suspensions from both strains were combined and plated on 25 cm x 25 cm large trays with LB agar supplemented with 0.4 mM DAP. For conjugation, plates were incubated overnight at 30°C. All cells were then harvested from mating plates by rinsing with PBS and plating on selection plates with LB agar supplemented with 100 µg/mL kanamycin, without DAP. After colonies of sufficient size appeared, transformants were harvested by scraping all cell mass from the plate and collecting the pooled scrapings in 1.5 mL tubes. The mutant library was diluted tenfold and then immediately frozen at -80°C. For competition experiments, a 1 mL 10-fold diluted aliquot (pool of all conjugations, ∼1 M CFU) was used to inoculate pre-cultures. The identification of mutant insertion sites (’transposon mapping’) is described in detail in Jahn et al., 2021.

### Growth medium

Strains were cultivated on complete (LB) medium, or minimal medium depending on experimental setup. Minimal medium was composed of 0.78 g/L NaH_2_PO_4_, 4.18 g/L Na_2_HPO_4_ × 2H_2_O, 1 g/L NH_4_Cl, 0.1 g/L K_2_SO_4_, 0.1 g/L MgCl_2_ × 6H_2_O, 1.6 mg/L FeCl_3_ × 6H_2_O, 0.4 mg/L CaCl_2_, 0.05 mg/L CoCl_2_ × 6H_2_O, 1.8 mg/L Na_2_MoO_4_ × 2H_2_O, 0.13 g/L Ni_2_SO_4_ × 6H_2_O, 0.07 mg/L CuCl_2_ × 2H_2_O. All components were added to autoclaved sodium phosphate buffer from filter-sterilized stock solutions. Different compounds were added as carbon and energy sources depending on the experimental setup (Table S1). Standard batch cultures were grown in a volume of 10 to 20 mL medium in 100 mL shake flasks at 30°C and 180 rpm. Precultures of the barcoded *C. necator* transposon library were supplemented with 200 µg/mL kanamycin and 50 µg/mL gentamicin to suppress growth of untransformed *C. necator* recipient or *E. coli* donor cells.

### Pressure bottle cultivation

Pressure bottles were used for batch cultivation in lithoautotrophic or nitrate respiration conditions. For lithoautotrophic growth, precultures of the *C. necator* transposon library were grown overnight in 10 mL LB medium, then transferred to 20 mL of minimal medium containing 2 g/L fructose. The precultures were used to inoculate 100 mL Duran Pressure Plus bottles containing 30 mL of minimal medium. Samples of the precultures were immediately frozen at -20°C for later use as T0 samples. A GB100 Plus gas mixer (MCQ instruments) paired with a GW-6400-2 anaerobe gas exchange system (GRinstruments) was used to flush and then fill the bottles to a final pressure of 1 bar above standard pressure, with a mixture of 70% H_2_, 15% O_2_ and 15% CO_2_. These culture bottles were shaken at 30°C, 150 rpm. When the cultures approached an OD_600_ of 1.0, they were diluted with minimal medium to OD_600_ of 0.1 and the gas replenished to ensure continuous growth occurred until the end point of 12 or 16 generations of growth were reached. Samples were taken at 4, 8, 12, and 16 generations. For nitrate respiration, the same protocol was used except that the minimal medium was supplemented with 2 g/L fructose and 1 g/L NaNO_3_. The headspace of bottles was flushed and then pressurized to 1 bar above standard pressure with 100% N_2_.

### Bioreactor cultivations

For chemostat experiments, the *C. necator* H16 transposon mutant library was grown as described previously (Jahn et al., 2021). Briefly, an 8-tube MC-1000-OD bioreactor (Photon System Instruments, Drasov, CZ) was customized to perform chemostat cultivation (Jahn et al., 2018, Yao et al., 2020). Bioreactors (65 mL) were filled with minimal medium supplemented with the respective carbon and nitrogen source, and inoculated with an overnight preculture to a target OD_720nm_ of 0.05. Bioreactors were bubbled with air at a rate of 12.5 mL/min and a temperature of 30°C. The OD_720nm_ and OD_680nm_ were measured every 15 min. For chemostat mode, fresh medium was continuously added using Reglo ICC precision peristaltic pumps (Ismatec, GER). For turbidostat mode, the target OD_720_ was set to 0.5 for fructose or 0.2 for formate cultures, and 4 mL medium (6.2% of total volume) was added as soon as the set point was reached. For library competition experiments, 15 mL samples were taken after 0, 8 and 16 generations of growth (population average). Cells were harvested by centrifugation for 10 min at 5,000 xg, 4°C, washed with 1 mL ice-cold PBS, transferred to a 1.5 mL tube, and centrifuged again for 2 min at 8,000 xg, 4°C. The supernatant was discarded and the pellet frozen at -20°C. For batch cultures in the bioreactor, WT or mutant strains were grown using the same setup but with 50 mL volume instead of 65 mL. The entire culture was harvested after the stationary phase was reached to determine the biomass yield.

### Biomass yield and growth rate

To determine biomass yield, 50 mL cell suspension was harvested by centrifugation for 10 min, 5,000 xg, 4°C. The pellet was washed twice with 1 mL mqH_2_O, transferred to preweighed 1.5 mL tubes and dried for 4 h at 70°C or 55°C overnight. Dried cell mass was measured on a precision scale and yield was calculated according to the equation: yield [gDCW / g substrate] = DCW [mg] / (substrate [g/L] * volume [L] * 1000). Growth rate was calculated by applying a sliding window of 5 hour length to the log OD_720_ over time, and fitting a linear model to each window. The slope of the fitted model was the growth rate for the respective window. From all slopes over time, the maximum was selected as the maximum growth rate.

### Hydrogen evolution experiment

Precultures of the H16 wild-type and the hydrogenase negative strain (Δ*hoxG*, Δ*hoxH*, Δ*hoxC*, Δ*hofG*) were grown in FN medium (Lenz et al., 2018) containing 0.4% fructose for 48 h at 37°C and 120 rpm. Main cultures were grown overnight in FGN medium (2 g/L fructose and 2 g/L glycerol) in baffled Erlenmeyer flasks (filled to 20% of the nominal volume) at 30°C and 120 rpm until an optical density at 436 nm (OD_436_) of approximately 6-8 was reached. The OD_436_ of the culture was adjusted to 6.5 using H16 buffer (C-free minimal medium). The cultures were transferred into 120 mL serum bottles (total bottle volume approx. 147 mL). The bottles were filled with 74 mL of the main culture and thus the gas space was 73 mL and the bottles were subsequently closed airtight with a rubber septum. The bottles were incubated at 30°C and 120 rpm and the gas composition was regularly analyzed by drawing 1 mL of gas from the bottles per measurement using a gas-tight 1-mL Hamilton syringe. The GC measurements followed published protocols (Schäfer et al., 2013).

### Gene fitness analysis (BarSeq)

Frozen cell pellets from the pulsed and continuous competition experiments were resuspended in 100 µL of 10 mM Tris and genomic DNA was extracted from 10 µL of the resuspension using a GeneJet Genomic DNA Purification Kit (ThermoScientific). Amplification of barcodes from genomic DNA was conducted using one of the custom forward indexing primers (BarSeq_F_i7, Table S2) and the reverse phasing primer (BarSeq_R_P2_UMI). For each sample 9 µL of genomic DNA extract (≥10 ng/µL) was combined with 3 µL of a forward indexing primer (100 nM), 3 µL of the reverse phasing primer pool (100 nM) and 15 µL of Q5 Mastermix (New England Biolabs). Cycle conditions were 4 minutes at 98°C followed by 20x (30 seconds at 98°C, 30 seconds at 68°C and 30 seconds at 72°C) with a final extension of 5 minutes at 72°C. Concentration of each sample was quantified using a Qubit dsDNA HS Assay Kit (Invitrogen). Samples were then pooled with 40 ng from up to 36 different samples being combined and run on a 1% agarose gel. Gel extraction was performed on the thick band centered around 200 bp and the concentration of the purified pooled library was quantified again via Qubit assay and diluted to 2 nM. The 2 nM library was then diluted, denatured and sequenced using a NextSeq 500/550 High Output Kit v2.5 (75 Cycles, Illumina) run on an Illumina NextSeq 550 instrument according to the manufacturer’s instructions. Library loading concentration was 1.8 pM with a 1% phiX spike. Calculation of gene fitness was performed using the rebar pipeline (https://github.com/m-jahn/rebar, Jahn et al., 2021), which trims and filters reads, extracts barcodes, and summarizes read counts per barcode. Fitness score calculation based on the log_2_ fold change of read count per barcode over time was implemented as an R script as part of the pipeline.

### Statistical analysis

Bioreactor cultivations, LC-MS/MS measurement for proteomics, and library competition experiments (’BarSeq’) were performed with four independent biological replicates. Here, biological replicate means that samples were obtained from independently replicated bottle or bioreactor cultures inoculated with the same preculture. If not otherwise indicated in figure legends, points and error bars represent the mean and standard deviation. No removal of outliers was performed. All analyses of fitness data, cultivations as well as proteomics results are documented in R markdown notebooks available at https://github.com/m-jahn/R-notebook-ralstonia-energy. A significance threshold of |f| ≥ 2 after at least 8 generations was chosen based on the bulk fitness distribution of mutants. Clustering of genes was performed based on similarity of fitness scores using the R function `hclust()` from package `stats` with method `ward.D2`. Optimal cluster number was determined using silhouette width analysis with the function `silhouette()` from R package `cluster`. T-SNE dimensionality reduction of genes was performed using R package `tsne`. Gene enrichment for KEGG pathways was performed by using the function `kegga()` from R package `limma`. The R statistical language was used with version 4.3.2.

### Data and software availability

Mass spectrometry proteomics data were obtained from the PRIDE repository with the dataset identifier PXD024819. Protein quantification results can be browsed and interactively analyzed using the web application available at https://m-jahn.shinyapps.io/ShinyProt. Sequencing data for BarSeq experiments are available at the European Nucleotide Archive with accession number PRJEB43757. The data for competition experiments performed with the transposon mutant library can be browsed and interactively analyzed using the web application available at https://m-jahn.shinyapps.io/ShinyLib/. All analyses of fitness data, cultivations as well as proteomics results are documented in R notebooks available at https://github.com/m-jahn/R-notebook-ralstonia-energy.

## Supporting information

Supplemental Items

## Acknowledgements

We like to thank Ute Hoffmann for support with data submission. This study was financially supported by the Swedish Research Council Vetenskapsrådet (Grant number 2016–06160), the Swedish Research Council Formas (Grant number 2015–939 and 2019–01491), and Novo Nordisk Fonden (Grant number NNF20OC0061469). Stefan Frielingsdorf and Oliver Lenz thank the cluster of excellence ‘UniSysCat’ under Germany’s Excellence Strategy EXC2008/1-390540038 for funding.

