## Supplemental Items for "The energy metabolism of *Cupriavidus necator* in different trophic conditions"

for the manuscript entitled:

### Supplementary Figures

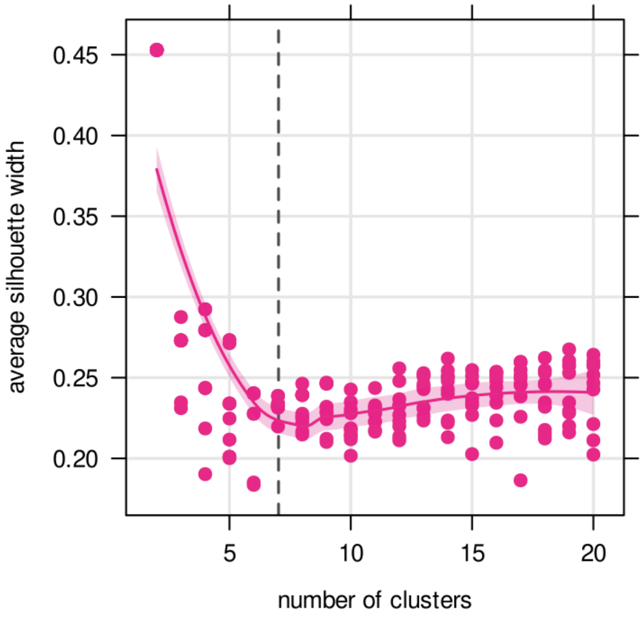


**Figure S1: Silhouette analysis for cluster separation.** To determine the optimal number of clusters, silhouette analysis was performed. The dataset was clustered from 2 to a maximum of 20 individual clusters, and the silhouette analysis was repeated 10 times for each number of clusters. Individual silhouettes (points) per cluster represent the quality of separation from their neighboring clusters. The average silhouette width is drawn as a smoothed line with error margins (95% confidence interval). A range of 7 to 20 clusters seemed to support roughly equally good separation. A number of 7 clusters was chosen as a good compromise between separation of all clusters and fragmenting the data into too many clusters (overfitting).

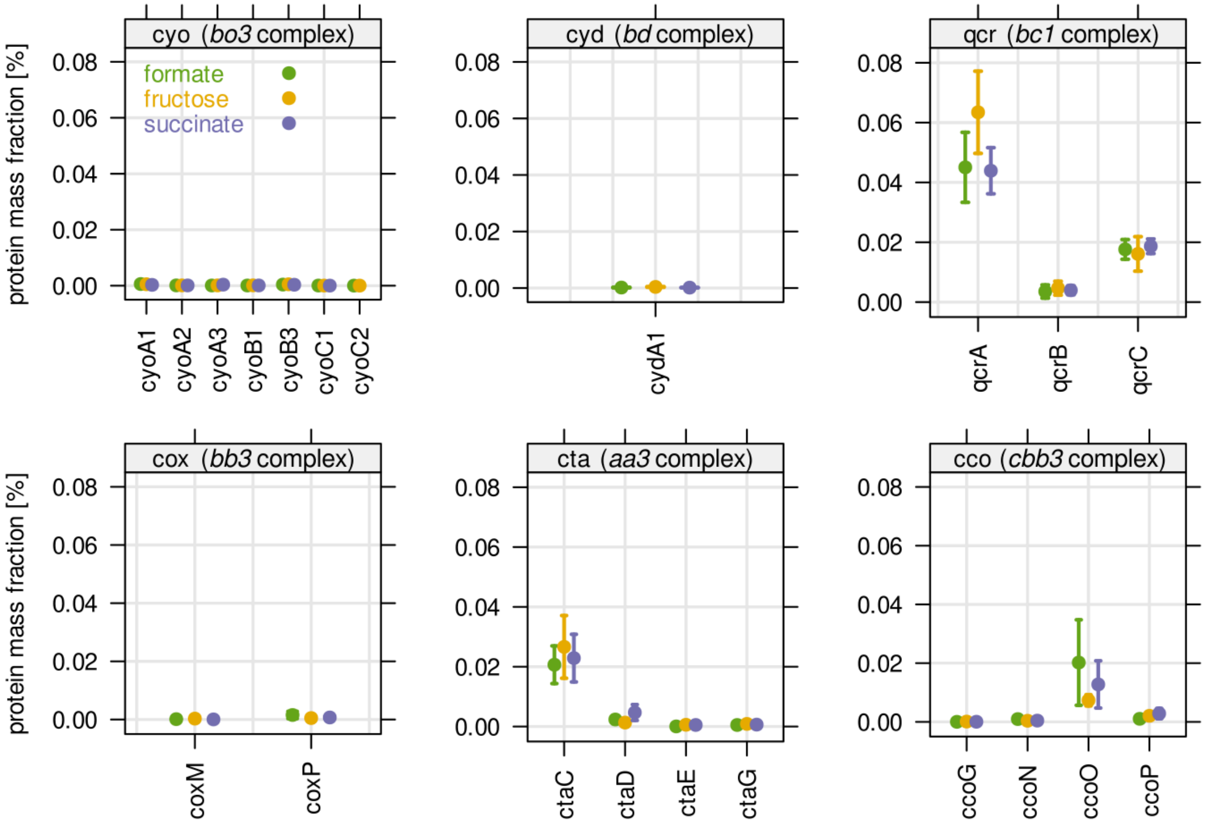


**Figure S2: Protein mass fraction for selected electron transport chain complexes.** Protein mass is given in % of the total proteome. Note that all of these complexes are located in or at the cytoplasmic membrane, and membrane proteins are often underrepresented in total proteome mass spectrometry because of their hydrophobic nature.

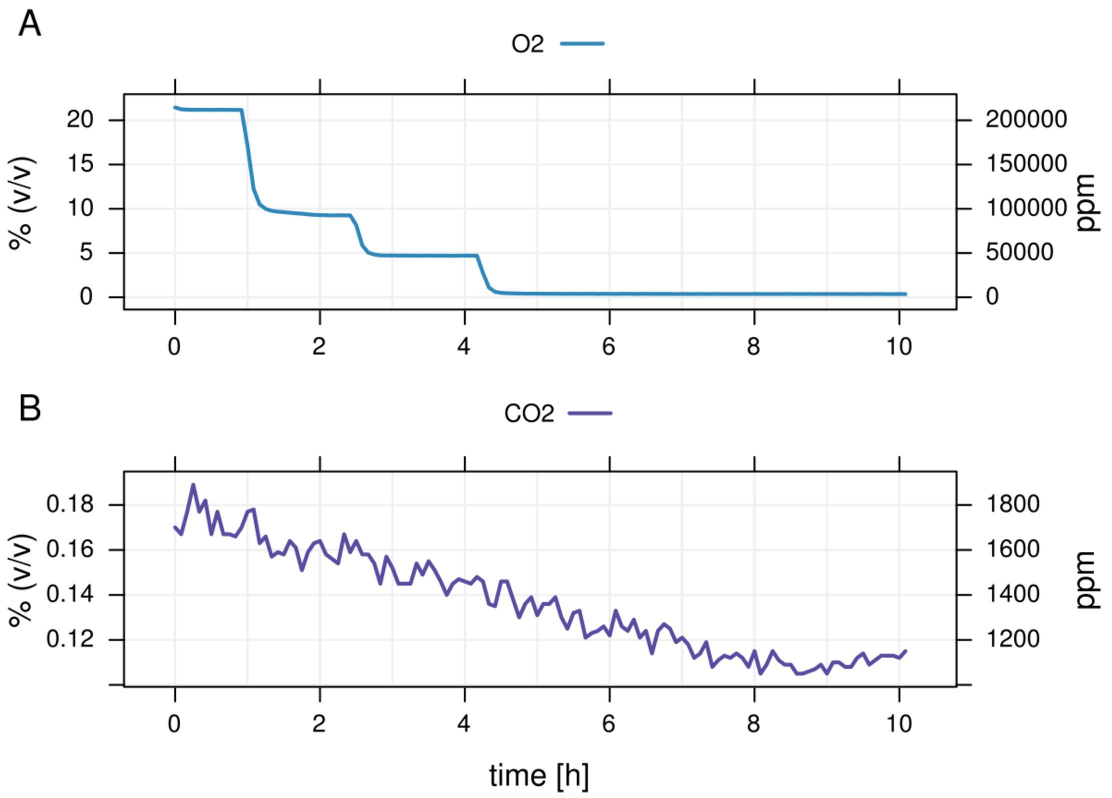


**Figure S3: Offgas analysis for nitrate respiration bioreactor experiments. A)** Oxygen ($O_2$) concentration for the first ten hours of a chemostat bioreactor cultivation, one selected replicate. *C. necator* H16 was grown in 65 mL tubes supplemented with 0.5 g/L fructose and 0.5 g/L $NaNO_3$. The nominal oxygen concentration was gradually lowered from 20% to 10%, 5% and 0% by mixing air and nitrogen ($N_2$) at a ratio of 50:50, 25:75 and 0:100, respectively. **B)** $CO_2$ concentration for the same time period as in A). $CO_2$ was emitted by the bacterial culture metabolizing fructose.

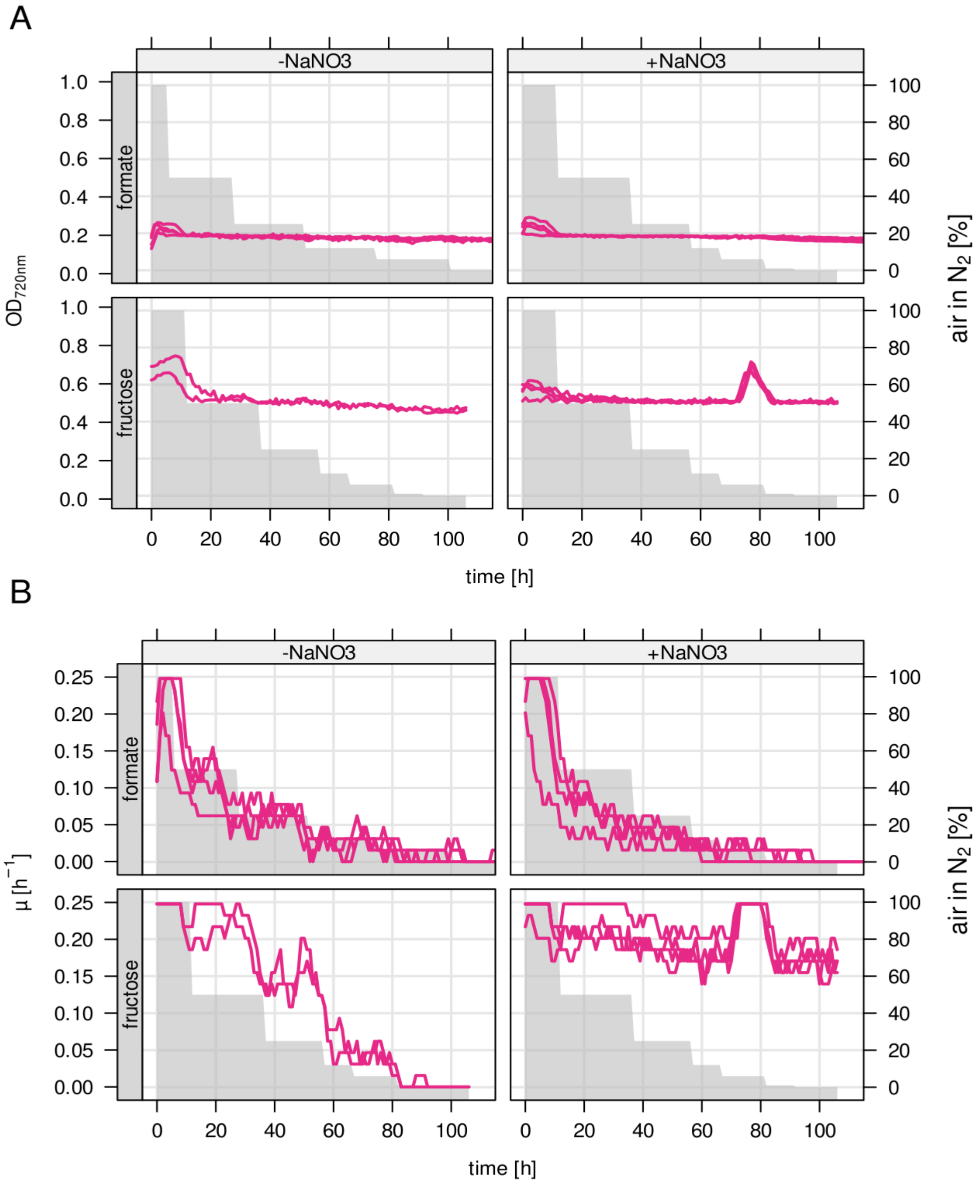


**Figure S4: Turbidostat bioreactor experiments to verify dependency on nitrate. A)** *C. necator* H16 was grown in 65 mL bioreactors in turbidostat mode. Minimal medium with 1.5 g/L formate (top row) or 0.5 g/L fructose (bottom row) was added at the same rate as the culture was growing, keeping the biomass density constant. Cultures were either supplemented with 1.5 g/L $NaNO_3$ as an alternative electron acceptor (right column), or not supplemented at all (left column). The oxygen concentration was gradually reduced from 20% to 0%. Red line and left axis, optical density at 720 nm. Grey area and right axis, % air in $N_2$. **B)** Growth rate μ determined for the same cultures (red line and left axis). Growth rate was calculated by counting the number of dilutions with a known volume of 4 mL/65 mL (6.2%) over time.

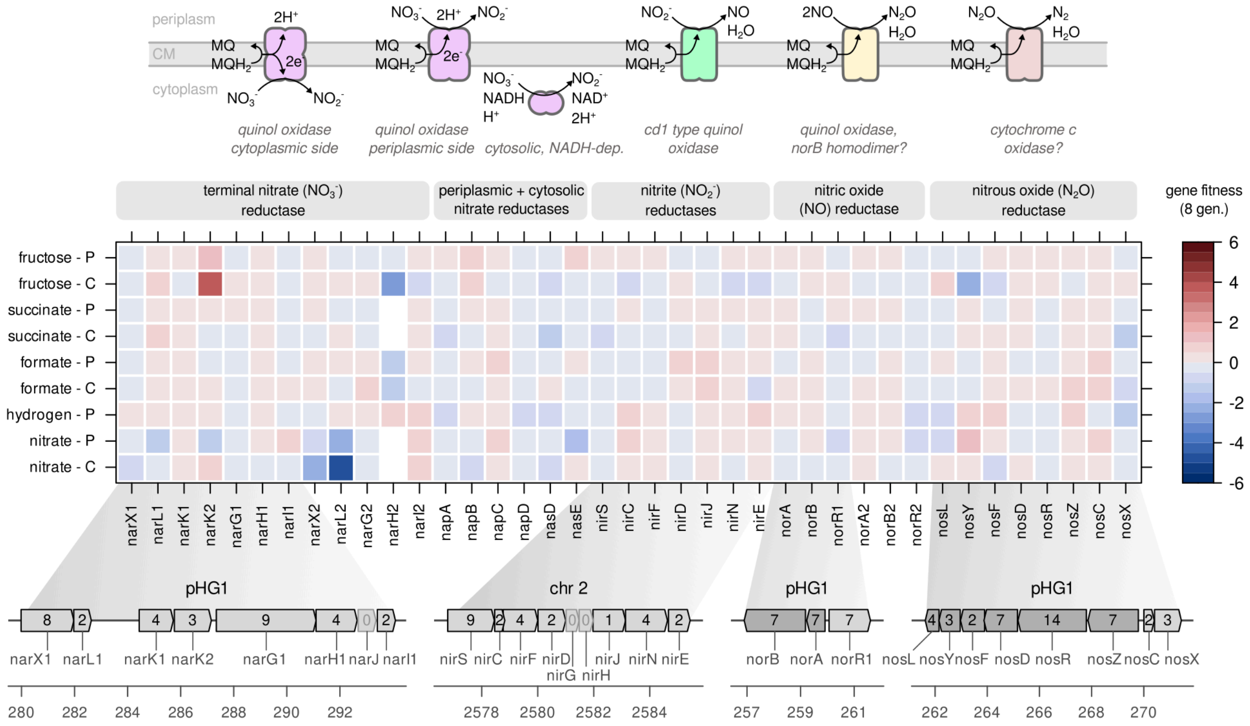


**Figure S5: Fitness for denitrification genes**. Fitness score for genes involved in one of the four steps of denitrification. The four major enzyme classes are nitrate ($NO_3^-$) reductases (purple), nitrite ($NO_2^-$) reductases (green), nitric oxide (NO) reductases (yellow), and nitrous oxide ($N_2O$) reductases (red). The respective enzymes and redox reactions are drawn schematically at the top of the figure. ***Nitrate reduction*** ($NO_3^- + QH_2 \rightarrow NO_2^- + Q + 2H^+$): The genome of *C. necator* contains three enzymes able to catalyze this reaction. The most important is the membrane-bound quinol oxidase (Kohlmann et al., 2014), encoded by two sets of *nar* genes (*narGHIJ*). Two additional operons encode structurally different nitrate reductases: *napABCD(E)* is located on pHG1 and encodes a membrane-bound quinol oxidase where nitrate reduction occurs at the periplasmic side of the membrane, and *nasD/E*, an assimilatory, soluble, cytoplasmic nitrate reductase located on chromosome 2. ***Nitrite reduction*** ($NO_2^- + QH_2 \rightarrow NO + H_2O$): Catalyzed by a *cd₁* type quinol oxidase encoded by the *nirSCFDGHJNE* operon on chromosome 2. For two very short genes, *nirG* and *nirH*, no transposon mutant was available. ***Nitric oxide reduction*** ($NO + QH_2 \rightarrow N_2O + H_2O + Q$): Catalyzed by nitric oxide reductase (*norAB* operon). *NorB* is presumably the catalytic subunit (assembles to a homodimer), and *norA* is a di-iron-center NO-binding protein with so far unknown function. Transcriptional regulator *norR* induces *norAB* expression in the presence of NO. Two copies of the operon and the regulator exist, one on pHG1 and one on chromosome 2. ***Nitrous oxide reduction*** ($N_2O + QH_2 \rightarrow N_2 + H_2O + Q$): Catalyzed by nitrous oxide reductase. The catalytic subunit is *nosZ* while *nosDFYL* presumably encode proteins that help with insertion of the copper cofactor into the functional enzyme. *NosR* is a transcriptional regulator. The scale bar indicates gene fitness scores after 8 generations of growth. Numbers inside genes represent the number of transposon mutants for that gene.

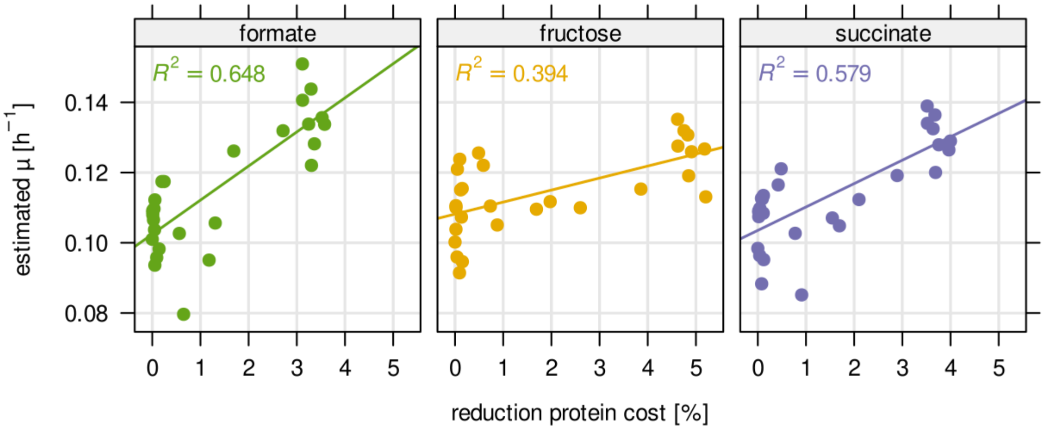


**Figure S6: Correlation of estimated growth rate µ and reduction of protein cost for hydrogenase mutants.** Points represent mutants of all *hox* and *hyp* genes shown in main Figure 6, A,B. Growth rate was estimated from $\log_2$ FC over time from library competition experiments (Methods). Protein cost was calculated as the % mass fraction of the total proteome, derived from mass spectrometry measurement. Line, linear regression model. $R^2$, coefficient of determination.

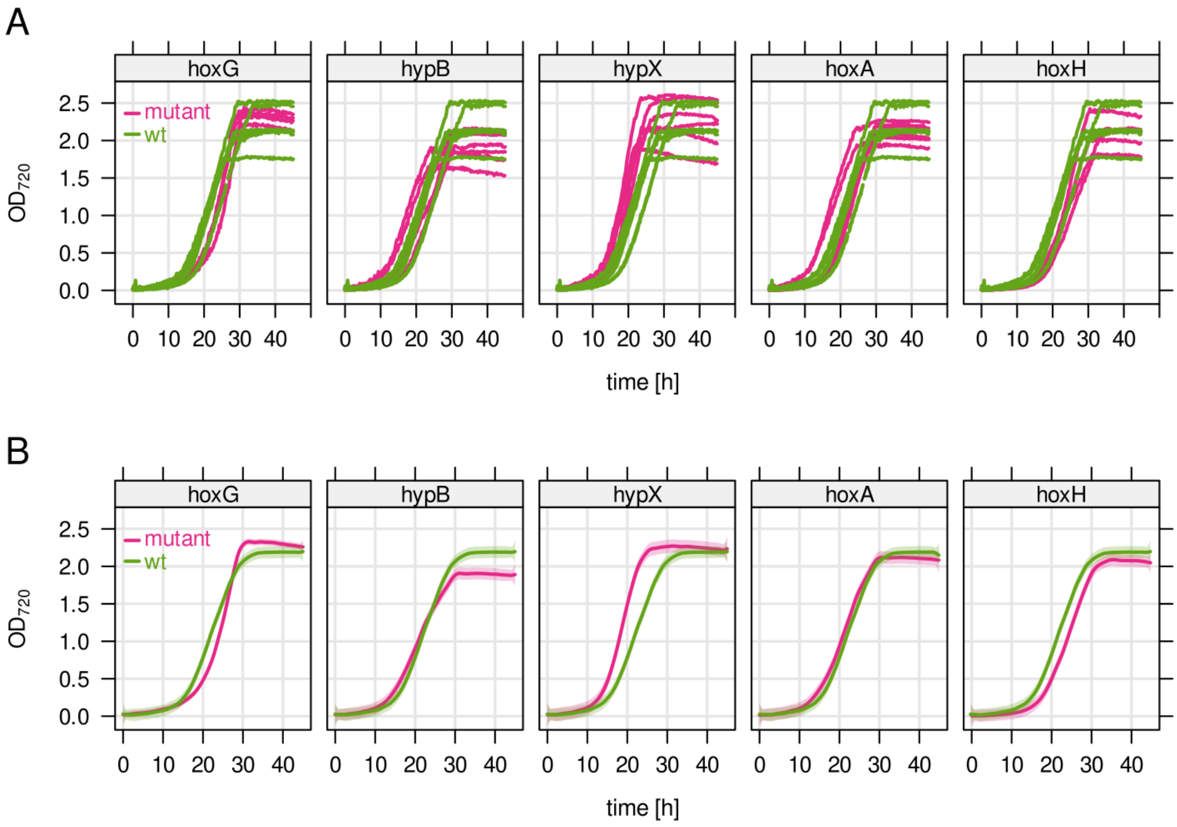


**Figure S7: Optical density measurement of *C. necator* hydrogenase mutants.** Batch bioreactor cultivation of in-frame deletion mutants for representative hydrogenase related genes, one mutant each for MH (*hoxG*) and SH (*hoxH*), two for accessory *hyp* genes (*hypB*, *hypX*), and the transcriptional regulator HoxA. Growth curves from the wild type control strain *C. necator* H16 are overlaid for comparison. Cultures were grown with 2.0 g/L fructose as the carbon and energy source, with at least 4 biological replicates. **B)** Smoothed average for the data in A). Semi-transparent ribbon: 95% confidence interval.

### Supplementary Tables

Table S1: Overview of *C. necator* transposon library cultivations for this study.

| **condition** | **feeding strategy** | **limiting substrate** | **sampling time [gen.]** | **reference** |
|---|---|---|---|---|
| Formatotrophy | chemostat, continuous | 1.5 g/L pH-neutralized formic acid | 0, 8, 16 | Jahn et al., 2021 |
| Formatotrophy | chemostat, 2h pulse | 1.5 g/L pH-neutralized formic acid | 0, 8, 16 | Jahn et al., 2021 |
| Heterotrophy, fructose | chemostat, continuous | 0.5 g/L D-fructose | 0, 8, 16 | Jahn et al., 2021 |
| Heterotrophy, fructose | chemostat, 2h pulse | 0.5 g/L D-fructose | 0, 8, 16 | Jahn et al., 2021 |
| Heterotrophy, succinate | chemostat, continuous | 0.5 g/L succinate | 0, 8, 16 | Jahn et al., 2021 |
| Heterotrophy, succinate | chemostat, 2h pulse | 0.5 g/L succinate | 0, 8, 16 | Jahn et al., 2021 |
| Nitrate respiration | chemostat, continuous | 0.5 g/L $NaNO_3$ | 0, 8, 16 | This study |
| Nitrate respiration | batch, repeated dilutions | 1.0 g/L $NaNO_3$ | 0, 4, 8, 12 | This study |
| Lithoautotrophy | batch, repeated dilutions | 70% $H_2$, 15% $O_2$, 15% $CO_2$ (1 bar) | 0, 4, 8, 12, 16 | This study |

Table S2: Oligonucleotides used in this study. Sequences with 'N' are variable portions of primers. Sequences in bold are non-homologous parts of primers such as restriction sites.

| # | name | type | sequence | reference |
|---|---|---|---|---|
| 1 | hofK270-XbaI_fwd | primer | **GCTCTAGAC**TCGTGCTGGTGCTTGAAGG | This study |
| 2 | PHG067-XbaI_rev | primer | **CATCTAGA**AGCTCGCATTGCGCGTAGTC | This study |
| 3 | pGE815 | plasmid | 3325bp XbaI-fragment (PCR) with 'hofK-hofG-PHG066' in pLO1 x XbaI | This study |
| 4 | pGE816 | plasmid | In frame deletion of a 1131bp SexAI-fragment of hofG in pGE815 | This study |
| 5 | BarSeq_F_i7 | primer | CAAGCAGAAGACGGCATACGAGAT NNNNNN GTGACTGGAGTTCAGACGTGTGCTCTTCCGATCTGATGTCCACGAGGTCTCT | Jahn et al., 2021 |
| 6 | BarSeq_R_P2_UMI | primer | AATGATACGGCGACCACCGAGATCTACACTCTTTCCCTACACGACGCTCTTCCGATCT NNNNNN GTCGACCTGCAGCGTACG | Jahn et al., 2021 |
